# A fitness landscape instability governs the morphological diversity of tip-growing cells

**DOI:** 10.1101/2023.06.12.544692

**Authors:** Maxim E. Ohairwe, Branka D. Živanović, Enrique R. Rojas

## Abstract

Cellular morphology affects many aspects of cellular and organismal physiology. This makes it challenging to understand the evolutionary basis for specific morphologies since the various facets of cellular physiology may exert competing selective pressures on this trait. The influence of these pressures, moreover, will depend on the mechanisms of cellular morphogenesis. To address this problem, we combined experimental cell biology with mechanics-based theory to analyze the morphological diversity of tip-growing cells from across the tree of life. We discovered that an instability in the convergent mechanism of “inflationary” growth shared by these cells leads directly to a bifurcation in their fitness landscape, which imposes a strong global constraint on their morphologies. Additionally, we found that co-selection for cell size and elongation rate explains variation among observable morphologies. This analysis rationalizes the morphology - and provides quantitative insight into the ecology - of an enormous diversity of important fungal, plant, protistan, and bacterial systems. Additionally, our study elucidates a fundamental principle of evolutionary-developmental biology that would be difficult to rigorously demonstrate in more complex systems.

## Introduction

It is often taken for granted that cellular morphology is functional [1], and yet in relatively few cases has the function or selective advantage of this trait been explicitly demonstrated [2, 3]. The comma shape of *Vibrio cholerae* cells, for one example, aids the “corkscrewing” of this bacterial pathogen into the host epithelium. In other systems, like fish keratocytes, morphology may be viewed as a consequence rather than a cause of cellular function. The highly variable crescent morphology of these cells emerges from the cytoskeleton dynamics that also drive cellular motility, yet motility is not dependent on a specific morphology [4]. Indeed, there are many examples of strong correlations between cellular morphology and cell state, type, or function [5, 6, 7], but these correlations do not necessarily mean that morphology serves a specific function.

On the contrary, cellular morphology affects *many* critical functions that cells perform. During fungal, plant, and animal development, for example, cellular morphology influences the assembly of the cytoskeleton [8, 9, 10], which governs myriad sub-cellular processes as well as the tissue morphogenesis that, in turn, feeds back onto cellular morphogenesis [11, 12]. Similarly, neuronal morphology affects action-potential propagation [13], neuronal connectivity [14], and signal integration [15]. Therefore, in general, cellular morphology is likely to be subject to many competing evolutionary pressures, whose effect will also be constrained by the mechanisms of cellular morphogenesis. Given this complexity, a major open challenge in biology is to identify and weigh the multiple factors that determine cellular morphology for a given system, that is, to understand cell morphology quantitatively as a complex trait.

Tip-growing cells (Fig. 1B,C) provide a simple example of the morphology-function question. These cells are found in a polyphyletic group of species that encompasses nearly all fungi, many important protistan pathogens, pollen tubes and root hairs in plants, as well as the phylum of bacteria that produces our best antibiotics. On one hand, it is clear that the filamentous morphology of tip-growing cells is intimately tied to their common function, which is to locate and consume nutrients (or deliver the gamete in the case of pollen tubes). On the other hand, the apical morphology of tip-growing cells is variable (Fig. 1A,S1A) and it is completely unknown if specific morphologies contribute to cellular function(s) or whether this variation simply reflects that in the underlying mechanisms of morphogenesis. This question is poignant since tip growth has arisen many times via convergent evolution.

**Fig. 1.**
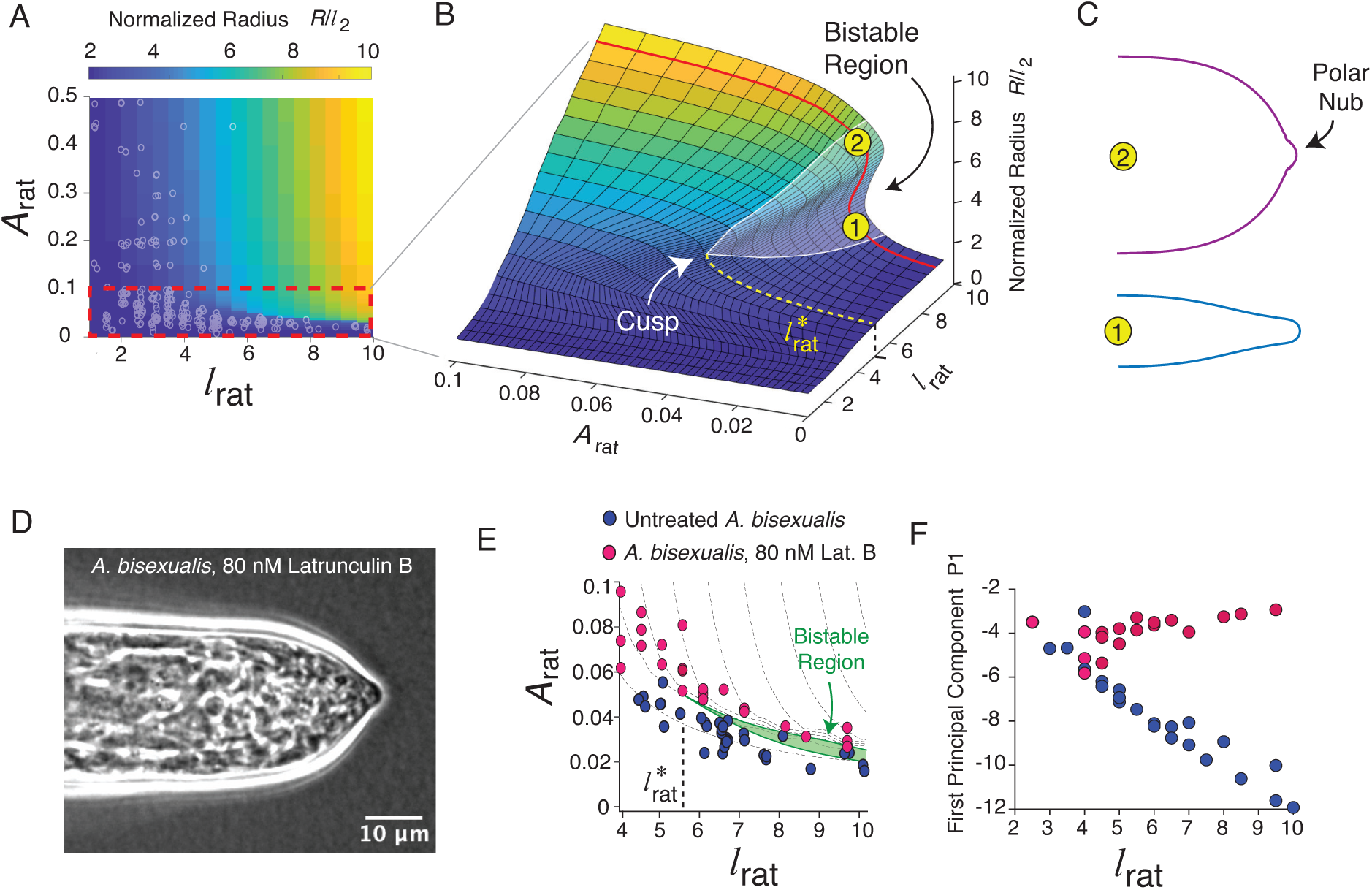
Tip-growing cells are found in diverse taxa. (A) Phase-contrast micrographs of three tip-growing cells representing protists (*Achlya bisexualis*), fungi (*Allomyces arbuscula*), and plants (*Lillium longiflorum*). (B) Time-lapse montage of an *Achlya bisexualis* hypha elongating. (C) The generic features of tip growth.

Evolutionary-developmental biology was conceived to address similar questions in animal systems, but the classic concepts and tools from this field have rarely been applied to single-cell morphology. A central tenet of evolutionary-developmental biology is that to understand morphological diversity across species, it is often sufficient to understand the mechanistic variation in their developmental programs. An important tool that emerged from this field was the “theoretical morphospace” [16], which is typically used to interpret global constraints on organismal morphology [17, 18]. In this light, using tip-growing cells as a model comparative system, our goal was to dissect the complex basis for cellular morphology by combining a mechanistic investigation of morphogenesis with a top-down analysis of morphological variation across taxa.

As for all walled cells, the morphology of tip-growing cells is defined by the geometry of the cell wall, which is a polysaccharide gel whose specific chemistry is variable across taxa (Fig. 1C). Because cell-wall geometry is determined by the apical cell-wall expansion that also leads to cell growth, morphogenesis and growth are the same process for these cells. In most cases, tip growth is self-similar such that apical morphology is approximately constant (Fig. 1B,2A). Therefore, the coordinated steady-state synthesis, metabolism, and physical deformation (expansion) of the apical cell wall are key processes underlying tip-growth morphogenesis. Tip-growing cells restrict cell-wall synthesis to the cell apex via localized exocytosis of cell-wall polysaccharides (Fig. 1C) [19, 20]. The actin cytoskeleton plays an important role in both exocytosis [21] and the spatial organization of the cell apex more broadly [22, 23]. How exocytosis is coordinated with cell-wall expansion is not fully understood. For several systems, however, it is clear that tip growth relies on a spatial gradient of cell wall biochemistry whereby the nascent, expanding cell wall is biochemically distinct from the mature, non-expanding wall (Fig. 1C) [24, 25, 26]. For example, during pollen tube growth, enzymes that prevent cell-wall cross-linking are included in the same exocytic vesicles that contain new cell-wall polysaccharides [27], ensuring that this material is mechanically soft [24]. Pollen tube growth, in turn, depends on the irreversible mechanical expansion of the soft apical cell wall by the turgor pressure within the cell [28]. Similarly, cell-wall expansion during the growth of fission yeast and the mating projections of budding yeast (examples of transient tip growth) is also driven by turgor pressure [29, 26]. In other words, tip growth is equivalent to controlled mechanical “inflation” of the cell for each of these systems.

**Fig. 2.**
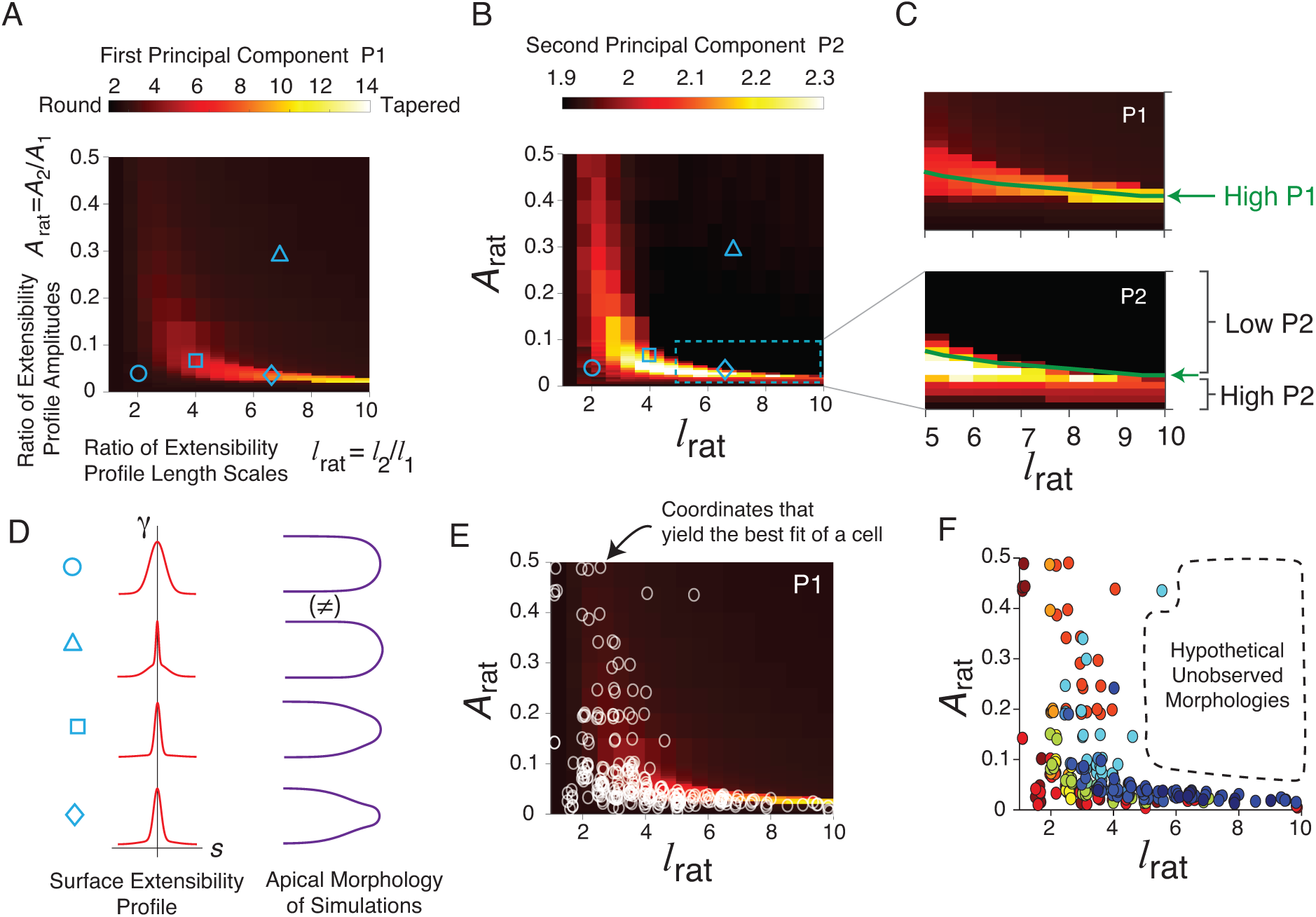
Tip-growing cells exhibit limited morphological variation. (A) Computationally extracted apical geometry versus time from a growing *A. bisexualis* hypha. (B) Key geometric variables of tip growth. (C-E) Time-averaged meridional curvature versus arclength for *A. bisexualis*, *A. arbuscula*, and *L. longiflorum* (red lines). Gray traces indicate the meridional curvature profiles from individual time points from which the averages were calculated. (inset) Reconstructions of the apical morphologies calculated the time-averaged meridional curvature profiles. The traces are representative of *n* = 32, 24, 32*and*32 cells, respectively. (F) The second principal variable (P2) versus the first principal variable (P1) for each cell from our multi-species principal component analysis data set (*n* = 9, 32, 36, 7, 32, 4, 10, 9, 23, 24, 32, 19 for *P.b.* sporangiophores, *A.b.*, *N.c.*, *P.b.* hyphae, *S.f.*, *M.t.*, *C.g.*,*S.v.*, *C.j.*, *A.a.*, *L.l.*, *S.p.*, respectively). (G) Apical geometry versus the first principal variable. For this calculation, the remaining principal variables were constant and equal to the average experimental value across all cells from all species. (H) Apical geometry versus the second principal variable, calculated as for P1.

Pioneering theoretical studies demonstrated that inflationary tip growth can, in principle, explain gross morphological variation of tip-growing cells from across nature [30, 31, 32]. Specifically, this mechanism can generate both round and tapered apical morphologies (Fig. 1A). However, inflationary tip growth was not tested in non-plant species, which are precisely those that exhibit tapered apical morphologies - it is possible that other mechanisms drive tip growth in these systems. Furthermore, the putative spatial relationship between cell-wall mechanical properties and cell-wall synthesis that generated morphological variability was not tested against cell biological data and was highly constrained. Accordingly, apical morphology relied on a single parameter (the theoretical morphospace was one-dimensional) and generation of tapered apical morphologies required a large spatial gradient in cell-wall thickness, which is not observed experimentally [33]. Therefore, it is unknown i) if diverse tip-growing systems use inflationary growth, ii) whether inflationary growth can generate natural morphologies in a manner consistent with underlying cell biology, and iii) whether tip-growing cells from across nature assume all morphologies realizable by inflationary growth.

By addressing these specific questions we identified strong mechanistic and evolutionary constraints on the morphological diversity of tip-growing cells from across nature. Through precise spatial measurements of the expansion rates and mechanical tensions in the cell walls of three divergent tip-growing systems - a protist, a fungus, and a plant - we first confirmed that each of them drives tip growth via inflation. We then systematically generalized previous theoretical models of inflationary growth [30, 31] to account for our data. This analysis revealed a generic mechanical requirement for the morphogenesis of prolate and tapered cells (Fig. 1A), and allowed us to describe the entire morphospace for tip-growing cells on mechanistic grounds. Surprisingly, we found that tip-growing morphologies from across nature populated a relatively small region of this morphospace. Further analysis revealed that an emergent cusp bifurcation [34] in the morphospace separated fast-growing natural morphologies from slow-growing hypothetical morphologies. We therefore interpret the morphospace as a fitness landscape and conclude that natural selection for fast growth imposes a strong constraint on the morphology of diverse tip-growing cells. Finally, we discovered that prolate and tapered morphologies (Fig. 1A) balance the benefits of cell size and elongation rate, which rationalizes the non-round morphology of most tip-growing systems. Collectively, these results explain the morphological variation of an enormous diversity of important cellular systems and provide a paradigmatic example of the interplay between cellular mechanics, cellular morphology, and natural selection.

## Results

### Tip-growing cells span a low-dimensional morphological space

To understand the morphological variation of tip growth, it was first useful to objectively quantify apical morphology across taxa. To do so, we recorded time-lapse phase-contrast micrographs (Fig. 1B, Movie S1) of hundreds of tip-growing cells from a broad diversity of organisms including fungi, protists, plants, and bacteria (Fig. 1A,S1A). We then computationally tracked cell-surface geometry from these micrographs (Fig. 2A). Position on the cell surface can be described by the co-ordinates *s*, the arclength from the cell pole, and *θ*, the azimuthal angle (Fig. 2B). We further quantified apical morphology by calculating the spatial profile of the curvature of a cell “meridian,” *κ*_s_(*s*) (Fig. 2B-E).

For all tip-growing cells, the meridional curvature is maximal at or near the cell pole (*s* = 0), and is zero at the cell equator, but the curvature profiles also revealed species-specific morphological signatures. For example, hyphae of the oomycete *Achlya bisexualis* exhibited a sharp curvature peak near the pole and a slight annular maximum distally, resulting in a tapered apical morphology (Fig. 2C). Hyphae of the chytrid fungus *Allomyces arbuscula* exhibited a curvature peak near the pole and annular curvature shoulders, resulting in a prolate morphology (Fig. 2D). As was reported previously, the maximum meridional curvature of *Lilium longiflorum* pollen tubes, *Medicago truncatula* root hairs, and *Schizosaccharomyces pombe* cells - each of which has round morphology - occurred in an annulus *≈* 1 *µ*m from the pole (Fig. 2E, S2B; [28, 35, 26]).

Despite these fine-scale features, when we applied principal components analysis to our entire multi-species data set, the first two principal variables (P1 and P2; Fig. 2F) accounted for 99.8% of morphological variation (Fig. S1C). Variation in the first principal variable corresponded precisely to scaling of apical morphology along the cell axis (Fig. 2G, S1D). As a result, P1 was also quantitatively correlated with the taper of the cell (Fig. S1E). The second principal variable was inversely related to the taper of the cell independent of axial scaling (Fig. 2H). Since taper is the most salient qualitative variable of tip-growing morphology (Fig. 1A), our objective quantification of morphological variation was consistent with our intuitive understanding of it and demonstrated that P1 and P2 are useful proxies for apical morphology. Moreover, variation of P1 and P2 alone explained both the annular maximum of pollen tubes and root hairs as well as the annular shoulders of fungal hyphae (Fig. S1F), revealing that these species-specific features are manifestations of low-dimensional geometrical variation.

### Turgor pressure drives tip growth in diverse organisms

Apical morphology is determined by the spatial dependence of cell-wall expansion across the cell apex (Fig. 3A). Therefore, we next quantitatively measured this dependence using *A. bisexualis*, *A. arbuscula*, and *L. longiflorum* as model systems since they largely encompass tip-growing morphological and phylogenetic diversity (Fig. 1A). To do so, we coated the cell wall with electrically charged fluorescent microspheres and recorded time-lapse phase-contrast and epifluorescence micrographs during tip growth (Fig. 3B,C, Movie S1,S2). The cell wall expands along both principal directions, *s* and *θ* (Fig. 3A). By computationally tracking the movement of the microspheres (Fig. 3D,S2A) and quantifying the rate at which they move away from each other on the cell surface (Fig. S2B-E), we calculated the principal expansion-rate profiles, *ε̇*_s_(*s*) and *ε̇_θ_*(*s*) (Fig. 3E-G; *Methods*, Eq. 5,6).

**Fig. 3.**
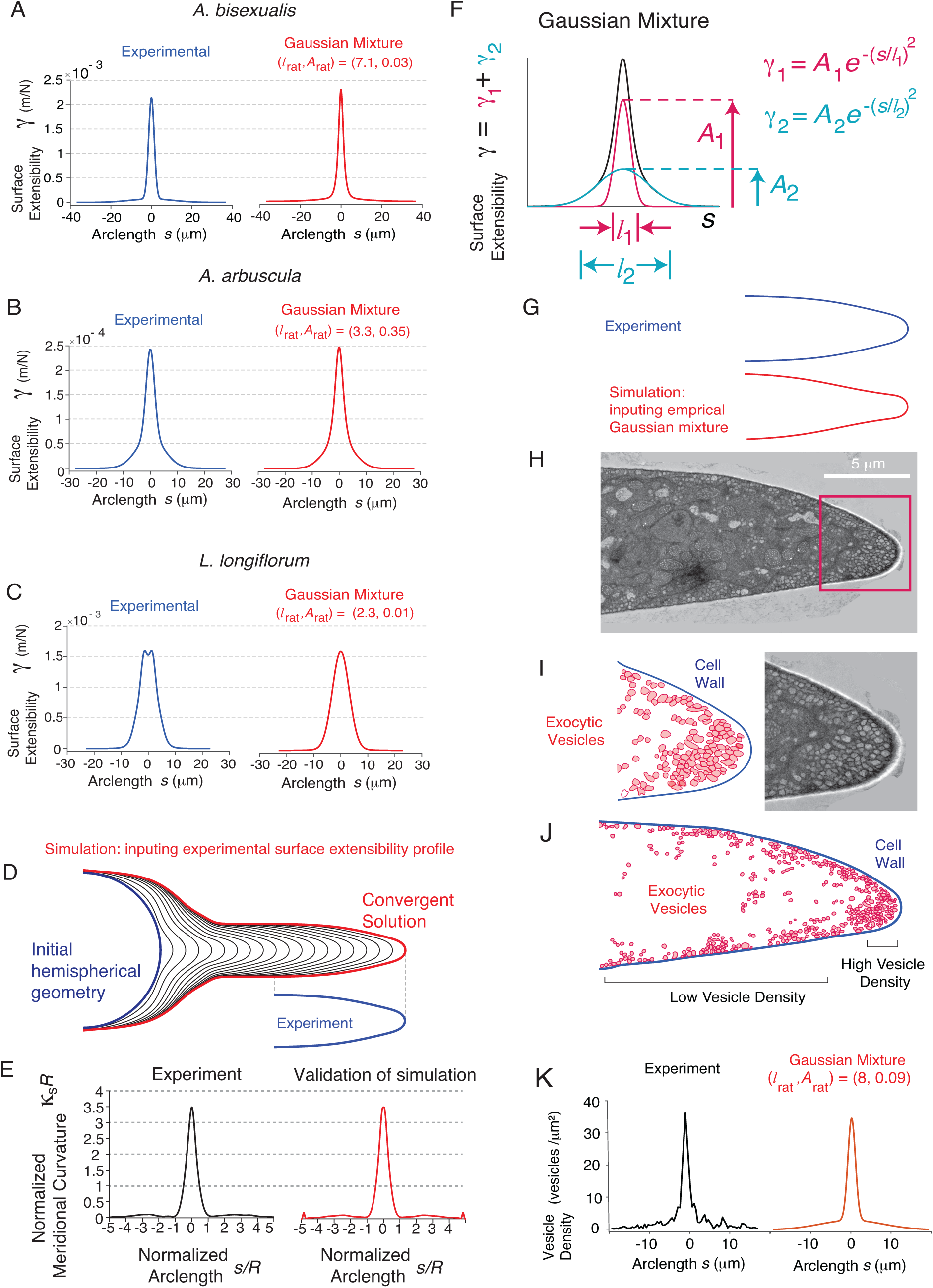
Diverse tip-growing species exhibit azimuthal expansion-rate anisotropy. (A) Key mechanical variables of tip growth. (B) Phase-contrast image of an *Achlya bisexualis* hypha. (C) Epifluorescence micrograph of the same hypha in (A) showing fluorescent microspheres attached to the cell wall. (D) Maximum intensity projection of the epifluoresence time-lapse micrograph from which the image in (C) was selected. This projection shows the paths of individual fluorescent microspheres in the lab frame of reference. (E-G) Principal cell-wall expansion rates versus arclength for *A. bisexualis*, *A. arbuscula*, and *L. longiflorum* cells (the data is representative of *n* = 2, 4, 4 cells, respectively). (H-J) Principal tensions versus arclength for the cells shown in (E-G). (K-M) Expansion-rate anisotropy versus tension anisotropy across the cell wall. Each trace represents a single cell. See main text for mathematical definitions of anisotropy.

Although the expansion-rate profiles must vary in order to generate distinct apical morphologies, the profiles from our three model systems all exhibited “azimuthal anisotropy” (*ε̇_θ_ > ε̇*_s_) except for at the pole (where they must be equal due to rotational symmetry) and the equator (where they are both zero). Previous low-resolution measurements of cell-wall expansion in several other tip-growing organisms [35, 36, 37, 38, 39, 26] were also consistent with azimuthal expansion-rate anisotropy, indicating that this is a widespread feature of tip growth.

To explore the consequences of azimuthal expansion-rate anisotropy on apical morphology, we developed a computational platform to simulate tip growth for a specific pair of principal expansion-rate profiles. We found that azimuthal expansion-rate anisotropy was necessary to generate tapered cells, but was not an intrinsic requirement for tip growth (Fig. S2F-L). It was previously pointed out that azimuthal expansion-rate anisotropy implies that turgor pressure drives cell-wall expansion since the principal tensions in the wall, which balance pressure, also exhibit azimuthal anisotropy (*λ_θ_ > λ*_s_) due to the generic filamentous morphology of tip-growing cells [35, 40]. To demonstrate this azimuthal tension anisotropy, we explicitly calculated the spatial profiles of the principal tensions from cell wall geometry (Fig. 3H-J; *Methods*, Eq. 9,10).

Turgor-driven “inflationary” growth has been demonstrated in several plant species [35, 41], as well as fission yeast [42], but not in other fungal or protistan systems. To explicitly test this mechanism, is not useful to directly compare the principal tension and expansion-rate profiles since the latter are also dependent on the mechanical properties of the cell wall, which are unknown and spatially inhomogeneous (Fig. 3A). To circumvent this issue, we hypothesized that the relative difference between the principal expansion rates will depend only on the relative difference between the principal tensions, regardless of the local mechanical properties. Therefore, we defined the expansion-rate and tension anisotropies quantitatively as: 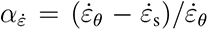 and *α_λ_* = (*λ_θ_ − λ*_s_)*/λ_θ_*. We found a precise monotonic scaling between these variables (Fig. 3K-M). Since there is no explanation for this dependence other than that wall expansion is caused by tension, this is strong evidence that inflation underlies tip growth across taxa.

### The spatial profile of cell-wall mechanical properties decays over two length-scales

Inflationary tip growth means that the spatial expansion-rate profiles (Fig. 3E-G), which lead to specific cell shapes (Fig. 2C-E), result from the spatial profiles of mechanical properties of the cell wall. While it is not possible to measure these properties directly, we inferred them by applying continuum-mechanics-based theory to our experimental data. A generic mechanical relationship between principal expansion rates and tensions is one in which they are related linearly [35]:

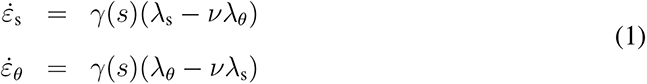

where *γ* and *ν* are mechanical properties. First, the “surface extensibility,” *γ*, determines the cell-wall expansion rate in a given direction if uniaxial tension is applied in that direction; this quantity accounts for both intrinsic mechanical properties (resulting from, for example, degree of cross-linking) and those in cell-wall thickness. Second, the “flow coupling,” *ν*, determines the rate at which the wall contracts in one direction due to uniaxial tension in the orthogonal direction. While the cell wall could possess non-linear mechanical properties for large or rapid deformations, this linear implementation of inflationary tip growth will be approximately valid at slow expansion rates characteristic of cell growth. It is unlikely, moreover, that higher-order corrections will meaningfully expand the morphospace since *γ*(*s*) is a powerful spatially dependent function. Finally, we found that our expansion-rate measurements are inconsistent with structural anisotropy in the cell wall (Fig. S2M), which could result from oriented synthesis of cell wall polysaccharides (*Methods*, Fig. S3), and that in any case such anisotropy would have a negligible affect on the morphospace that inflationary growth yields (Fig. S4). Indeed, our linear isotropic implementation of inflationary tip growth (Eq. 1) provided accurate, simultaneous fits of the experimental curvature (Fig. S5) and expansion-rate profiles (Fig. S2C-E; *Methods*) from each cell of each of our model species. Therefore, this theoretical formulation of tip growth is a useful tool for exploring its morphospace.

To this end, we calculated the experimental surface extensibility profiles, *γ*(*s*), for each model species by substituting their experimental principal expansion-rate and tension profiles into Eq. 1 (Fig. 4A-C). We found that these profiles possessed species-specific features that qualitatively resembled the curvature profiles. In particular, the profiles from *A. bisexualis* and *A. arbuscula* displayed sharp central peaks while the *L. longiflorum* meridional profile had a slight annular maximum. Both the *A. bisexualis* and *A. arbuscula* azimuthal profiles had clear “shoulders” that indicated that these profiles decayed to zero over two distinct length scales. We adapted our computational platform to simulate tip growth for a given extensibility profile. Inputing our experimental profiles generated apical morphologies that were indistinguishable from the experimental morphologies (Fig. 4D,E,S5), further validating the use of the linear implementation of inflationary tip growth (Eq. 1).

**Fig. 4.**
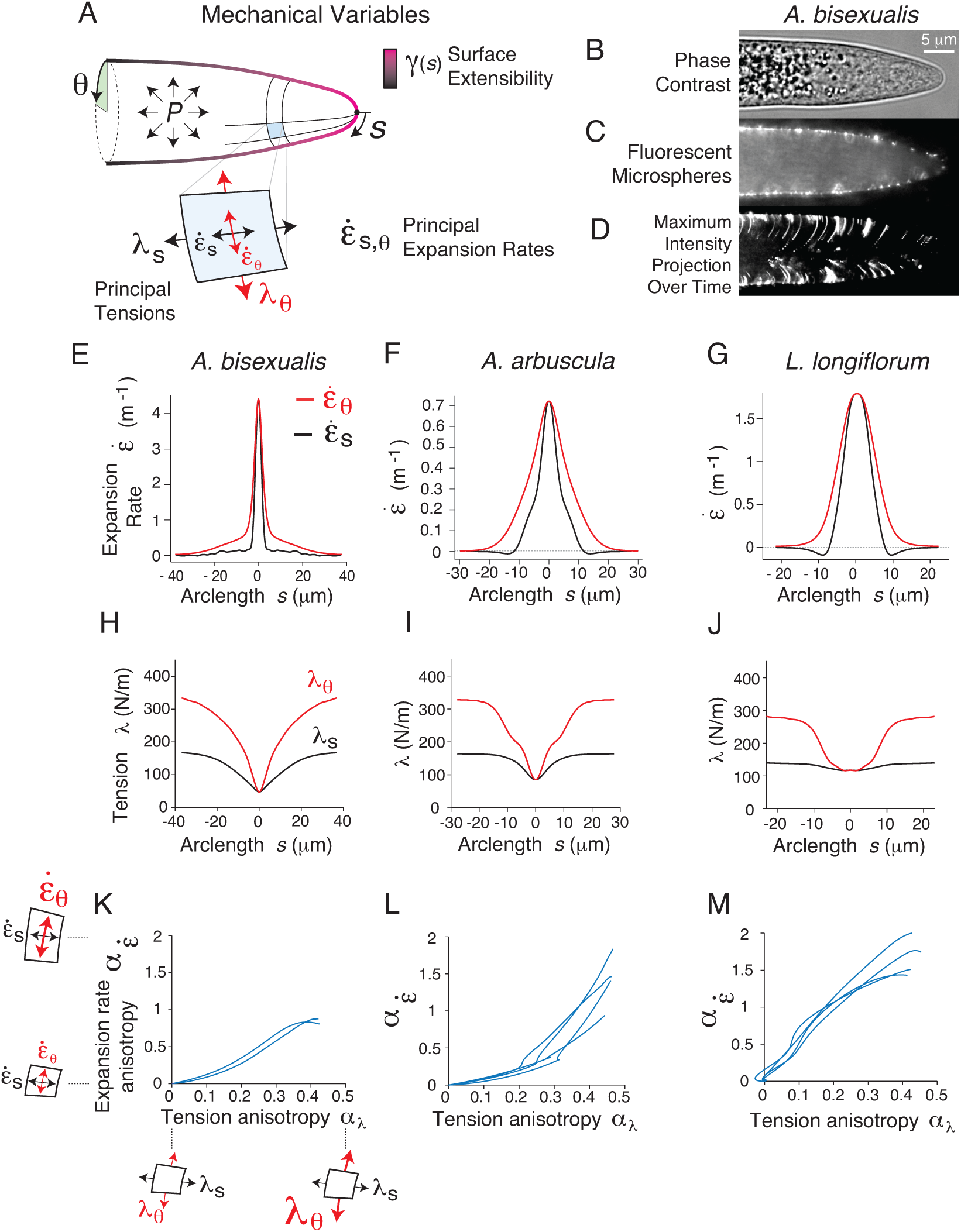
A two-parameter surface extensibility profile accounts for morphological diversity among tip-growing cells. (A-C) (left) Experimentally determined surface extensibility profiles for representative cells of *A. bisexualis*, *A. arbuscula*, and *L. longiflorum*. Similar results were obtained for *n* = 2, 4, 4 cells, respectively. (right) Best fit of the experimental surface extensibility profiles by an empirical Gaussian mixture function. (D) Computational simulation of tip growth based on Eq. 1 of the main text, inputting the experimental surface extensibility profile shown in (A), and beginning from a hemispherical initial condition. The simulation converged precisely to the experimental apical morphology. (E) (left) The normalized experimental meridional curvature versus normalized arclength. (right) The normalized meridional curvature of the simulation shown in (D) versus normalized arclength. (F) The empirical Gaussian mixture function used to fit the surface extensibility profiles. (G) Comparison of the experimental time-averaged apical morphology of an *A. bisexualis* hypha and the apical morphology of a computational simulation generated with an empirical Gaussian mixture surface extensibility profile shown in (A). (H) Transmission electron micrograph of a thin section of the mid-plane of an *A. bisexualis* hypha. (I) (left) Location of exocytic vesicles and cell wall in red box in (H). Zoom of the red boxed region in (H). (J) The location of the exocytic vesicles and cell wall in (H). (K) (left) The density of vesicles per unit cell wall length versus arclength. Similar results were obtained for *n* = 3 cells (Fig. S6A). (right) Best fit of the experimental vesicle density profile by the empirical Gaussian mixture function.

We hypothesized that we could accurately fit the surface extensibility profiles with a generic spatial function that was peaked at the cell pole and decayed over two length scales. We first tested a “Gaussian mixture” function (the superposition of two Gaussians; Fig. 4F). When the widths (*l*_1_*, l*_2_) and amplitudes (*A*_1_*, A*_2_) of the component Gaussians were used as fitting parameters, this function provided an accurate fit of the experimental profiles (Fig. 4A-C). To assess whether these empirical fits could generate realistic apical morphologies, we input them into our simulation platform. In each case, the simulations reproduced accurate apical morphologies: for *A. bisexualis*, the apical geometry of the simulation deviated slightly from the experimental geometry but accurately reproduced the tapered morphology (Fig. 4G,S5A), while for *A. arbuscula* and *L. longiflorum* the difference between the simulated and experimental morphologies were only distinguishable by minute differences in their meridional curvature profiles (Fig. S5B,C). Although a single Gaussian function provided an accurate fit of the *L. longiflorum* apical morphology (Fig. S5C), it provided a poor prediction of *A. bisexualis* and *A. arbuscula* morphology, demonstrating the requirement of a two-length-scale surface extensibility profile to generate tapered and prolate apical morphologies. We confirmed that these results did not depend on the specific functional form of the surface extensibility profile by simulating tip growth with non-Gaussian versions (Fig. S5).

In light of our analysis, we can summarize a mechanism of morphogenesis of prolate and tapered (i.e., non-round) apical morphologies. When subject to inflation by turgor pressure, a two-length scale surface extensibility profile, in combination with the intrinsic azimuthal anisotropy of the principal tension profiles, generates principal expansion rate profiles that correspond to roughly isotropic expansion near the pole and gradual azimuthal expansion in the sub-polar region (Fig. 3E,F). Such a two-length-scale surface extensibility profile is necessary and sufficient to generate even mildly tapered apical morphologies, which are characteristic of nearly all tip-growing cells.

### Exocytic vesicle density predicts cell wall mechanical properties

We next sought to identify the ultrastructural and/or biochemical factors that underlie the two length scales associated with the surface extensibility in protists and fungi. First, we used transmission electron microscopy to image thin cross-sections of fixed *A. bisexualis* hyphae, which have a highly tapered morphology. In contrast to the predictions of previous theoretical studies, we did not observe a strong gradient in cell-wall thickness (Fig. 4H). However, there was a clear spatial dependence of exocytic vesicle distribution within the cytoplasm: near the pole there was a large pool of vesicles whereas in the subpolar region there was a thin layer of cortical vesicles (Fig. 4H-J). The quantitative spatial dependence of vesicle density showed two clear length scales, whose values were consistent with the length scales associated with the surface extensibility profile (Fig. 4K,S6A). This supports a model in which exocytosis directly softens the cell wall.

Next, we tested the hypothesis that the two length scales in surface extensibility correspond to two biochemically distinct sections of the cell wall. To do so, we labeled the cell walls of *A. bisexualis* hyphae with the dye Direct Red 23, which preferentially binds to cellulose, and Calcofluor White, which non-specifically binds to various *β*-glucans [43]. We found that Direct Red 23 labeling was nearly uniform across the cell apex (Fig. S6B,C), indicating that variation in cellulose content does not mediate that in wall extensibility. Conversely, Calcofluor White labeling was absent in the apical section of the cell wall, but increased sharply in the subapical section (Fig. S6D-F). This suggests that synthesis of Calcofluor-binding *β*-glucan(s) may contribute to the determination of the longer of the two length scales associated with the surface extensibility profile.

### Diverse tip-growing cells exhibit a limited range of possible apical morphologies

The identification of two clear length scales in the surface extensibility profiles of our non-round model systems motivated us to test whether such a profile could accurately explain the natural apical morphologies across our comprehensive multi-species data set (Fig. 2F), and also to explore the entire theoretical morphospace that a two-length-scale profile could generate. Given such a surface extensibility profile, there are only three scalar parameters that determine apical morphology: *ν*, *l*_rat_ = *l*_2_*/l*_1_, *A*_rat_ = *A*_2_*/A*_1_, where *l_i_* and *A_i_* are the length scales and amplitudes of the component surface extensibility profiles (Fig. 4F; *Methods*). We simulated tip growth across these parameters and created a theoretical morphospace by calculating the dependence of the principal variables (from our multi-species PCA analysis) across this space (Fig. 5A-C). For a given flow coupling, most of the morphospace spanned by *l*_rat_ and *A*_rat_ had low values of P1, generating round apical morphologies (Fig. 5A,D). However, there was a narrow region of the morphospace that displayed high values of P1, corresponding to tapered morphologies. Importantly, although P1 is a useful proxy for cell shape, it did not completely determine apical morphology. In particular, the region of high P1 (tapered cells) separated regions of low P1 (rounder cells) that had different values of P2 (Fig. 5C). We found this morphospace provided an excellent fit of nearly every cell from every species (Fig. S7A,B).

Variations in the flow coupling, *ν*, shifted the region of tapered morphologies, but not the qualitative dependence of P1 on *l*_rat_ and *A*_rat_ (Fig. S7D). Away from the tapered region, for a given parameter coordinate, flow coupling had little effect on cell morphology (Fig. S7E). Furthermore, altering *ν* had no effect on the fitting power of the model across species (Fig. S7B). Thus, while for most systems we do not have information about the value of the flow coupling, we conclude that this parameter is not important for shape determination, but rather that *l*_rat_ and *A*_rat_ are the key morphogenetic parameters.

**Fig. 5.**
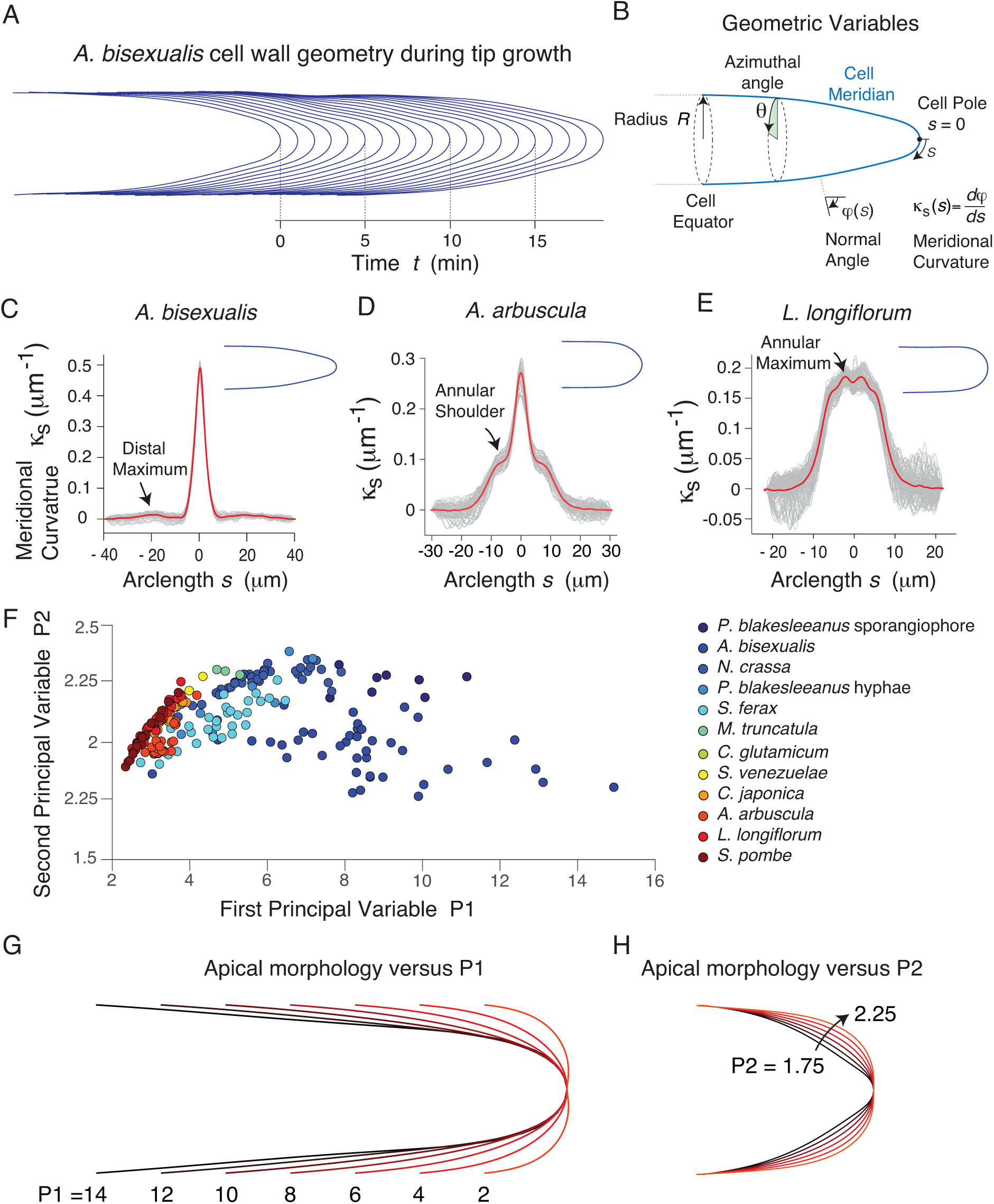
Tip-growing cells exhibit a limited range of possible morphologies. (A) The empirical Gaussian mixture surface extensibility profile. (A) The empirical Gaussian mixturefunction. (B) The first principal variable (P1) of simulations of inflationary tip growth (Eq. 1) inputting an empirical Gaussian mixture surface extensibility profile, versus *l*_rat_ and *A*_rat_. A flow coupling of *ν* = 0.5 was used. P1 is the first principal component of the PCA analysis of our multi-species data set. (C) The second principal variable (P2) of simulations of inflationary tip growth, versus of *l*_rat_ and *A*_rat_. (D) Apical morphologies at 4 points of the morphospace shown in (B) and (C). (E) Coordinates of (*l*_rat_*, A*_rat_) that, when used to simulate cell growth, yielded the best fit of the geometry for each cell of each species (white circles) overlaid onto the morphospace shown in (B). (E) Coordinates of (*l*_rat_*, A*_rat_) that, when used to simulate cell growth, yielded the best fit of the geometry for each cell of each species, color-coded by species.

Interestingly, we found that the values of *l*_rat_ and *A*_rat_ that provided the best fits for the cells in our data set populated a limited region of the morphospace, corresponding to a low value of one or both of these parameters (Fig. 5E,F). Furthermore, these (*l*_rat_,*A*_rat_) coordinates were sharply bounded by the region of tapered morphologies. In order to gain insight into this empirical constraint, we analyzed other metrics of cell geometry as a function of *l*_rat_ and *A*_rat_. We discovered that for a given value of *l*_1_ (the primary length scale in the the surface extensibility profile; Fig. 4F), that cell radius varied as a function of *l*_rat_ and *A*_rat_, with lower values of these variables yielding thinner cells (Fig. 6A). Given that low values of *l*_rat_ and *A*_rat_ also provided the best fits to experimental apical morphologies (Fig. 5E,F), this means that cells from across nature are relatively thin compared to the primary length scale of their surface extensibility profile. While the transition between thin cells to wide ones was gradual as *l*_rat_ was increased, it was extremely sharp as *A*_rat_ was increased for *l*_rat_ ≳ 5 (Fig. 6A).

**Fig. 6.**
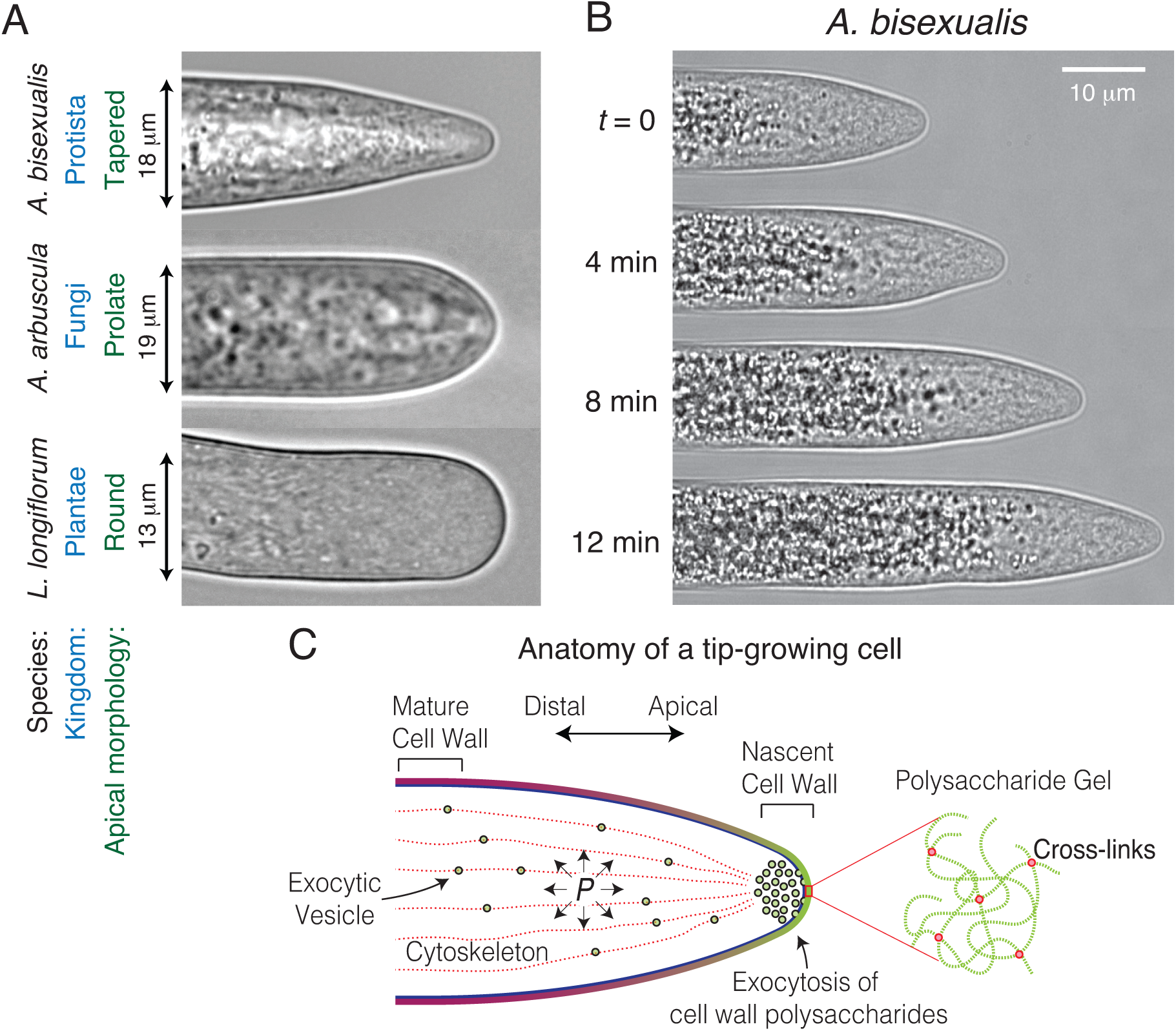
A cusp bifurcation instability constrains natural tip-growing cell morphologies. (A) Radius of simulated cells, normalized by *l*_2_, as a function of *l*_rat_ and *A*_rat_. The white circles indicate the same coordinates that yielded the best fit of the experimental morphologies shown in Fig. 5E,F. (B) Radius of simulated cells, normalized by *l*_2_, versus *l*_rat_ and *A*_rat_, within the region enclosed in the red dotted box in (A). The white shaded region indicates the bistable region of the morphospace. *l^∗^* is the value of *l*_rat_ above which bistability occurs. (C) The stable steady-state apical morphologies of simulated cells using a Gaussian mixture surface extensibility profile with (*l*_rat_*, A*_rat_) = (9.5, 0.025); the only difference between the simulations was the initial condition. The coordinates of the solutions are indicated by the yellow points in (B). (D) Phase-contrast micrograph of a *A. bisexualis* hypha treated with 80 nM latrunculin B. (E) Values of (*l*_rat_*, A*_rat_) that, when used to simulate cell growth, yielded the best fit of the apical geometry for *A. bisexualis* hyphae, comparing untreated hyphae to those treated with 80 nM latrunculin B. (F) The first principal component of the apical morphology of untreated and latrunculin-B-treated *A. bisexualis* hyphae versus *l*_rat_.

### A cusp bifurcation in the inflationary mechanism of tip growth constrains apical morphology

Sharp changes in the solutions of dynamical systems upon small changes in their parameters often reflect bistability. To explore this possibility, we performed a more fine-scale computational analysis of the region of the morphospace where cell radius changed sharply (Fig. 6A, red dotted box). As suspected, the sharp change in cell radius corresponded to the bistable region of a cusp bifurcation or “catastrophe” [34] (Fig. 6B). That is, above a critical value 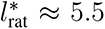, the manifold of model solutions folds such that for a narrow range of *A*_rat_ there exist two stable apical morphologies for a single surface extensibility profile. One of these morphologies is tapered, whereas the other is round but with a nub at the cell pole (Fig. 6C). Within this region, the apical morphology on which simulations converged depended on the initial condition (Fig. 6C,S7F). Accordingly, for values of 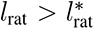, the system exhibited hysteresis when *A*_rat_ was cycled (Fig. S7F). Whereas many tip-growing species were confined to the low-radius region of the morphospace by virtue of having low values of *l*_rat_, species with tapered apical morphologies (*e.g.*, *A. bisexualis* and *P. blaakeseanus* sporangiophores) clustered below the bistable region, corresponding to low values of *A*_rat_ (Fig. 6A,S7G).

The bistable region in the morphospace results in what is colloquially known as a “tipping point” [44] with respect to changes in *A*_rat_. Intuitively, this tipping point arises due to competition between the two component surface extensibility profiles that comprise the net profile (Fig. 4F). The thinner of the two profiles, on its own, would generate a round cell with low radius, while the wider profile would generate a round cell with a large radius. In the limit that the amplitude of one of these profiles is much greater than the other, the profile with greater amplitude generates a round cell that grows much faster than the profile with lower amplitude, resulting in a round cell of either low or high radius. On the other hand, for a range of *A*_rat_, the two profiles generate cells that grow at similar rates, and therefore competition between these component profiles can generate tapered apical morphologies. However, tapered morphologies are nearly unstable because turgor pressure is directionless and will tend to “inflate” the conical sub-polar region of the cell. Finally, if the component profiles have similar widths (i.e., if *l*_rat_ is low) they will generate rounded cells of similar radii. In this case, it does not matter which solution grows faster, and the instability disappears at the critical point, or “cusp” (Fig. 6B).

Our analysis made the strong prediction that altering the surface extensibility profile of tip-growing cells would lead to previously unobserved apical morphologies. Specifically, because tapered morphologies are predicted to be nearly unstable, we hypothesized that mild nonspecific perturbations could induce them to adopt a round shape with a polar nub (Fig. 6C). To test this, we treated *A. bisexualis* hyphae with sub-inhibitory concentrations of latrunculin B, which inhibits actin polymerization and therefore interferes with the spatial organization of tip growth. Remarkably, this treatment resulted in stably growing hyphae with apical morphologies qualitatively similar to those predicted by our theory (Fig. 6D, Movie S3). These non-natural morphologies populated a region of the morphospace with slightly higher values f *A*_rat_ than that populated by untreated cells, and hyphae with *l*_rat_ values greater than *l*_rat_*^∗^* now clustered above the bistable region (Fig. 6E). Therefore, although latrunculin-B-treated cells cluster with untreated cells in the space spanned by *l*_rat_ and *A*_rat_ as would be expected from a mild perturbation, the apical morphologies of the two populations diverge because they reside on different branches of the manifold of stable apical morphologies. This is clearly illustrated by comparing the values of P1 as a function of *l*_rat_ for the two populations (Fig. 6F). Finally, latrunculin-B-treated cells were wider than untreated cells, in quantitative agreement with our theory (Fig. S7H).

To sum, small perturbations in key morphogenetic variables via sub-inhibitory pharmacological perturbation caused dramatic changes in cellular morphology, a hallmark of bistability.

The sharp morphological transition was precisely predicted by the inflationary model of growth with a two-length scale surface extensibility profile, and there is no reason to expect this transition otherwise. These results are further validation of the (linear isotropic) inflationary mechanism of tip growth and are strong evidence that the emergent cusp bifurcation associated with this mechanism imposes a strict, ubiquitous constraint on apical morphology.

### The morphospace of tip growth corresponds to a fitness landscape

Although the cusp bifurcation emerges mechanistically from inflationary growth, the discovery that it imposes a constraint on cellular morphology was empirical. To understand the basis for this constraint we next asked how other aspects of cellular physiology depended on morphology. Strikingly, we found that the rate of cellular elongation (normalized to the rate of total cell volume enlargement, *v/ρ*) varied strongly across the morphospace (Fig. 7A). Specifically, elongation rate was inversely related to cell radius. This result has a simple intuitive explanation: that for a given rate of cell-wall synthesis, a cell can either be thin and elongate rapidly or be wide and elongate slowly. This analysis made the clear prediction that if we induce tapered hyphae to adopt round, nubbed morphologies, that these wide cells would elongate more slowly. Indeed, mild treatments of latrunculin B that caused cells to adopt non-natural wide shapes also caused them to grow slower, in quantitative agreement with our theory (Fig. 7B). The observation that naturally occurring apical morphologies correspond to thin, fast-growing cells - and that hypothetical wide, slow-growing morphologies are mechanistically possible but not observed in nature - strongly suggests that the dependence of elongation rate on the morphogenetic parameters represents a fitness landscape, and that variation in apical morphology of tip-growing systems across the tree of life primarily reflects selection for fast elongation.

**Fig. 7.**
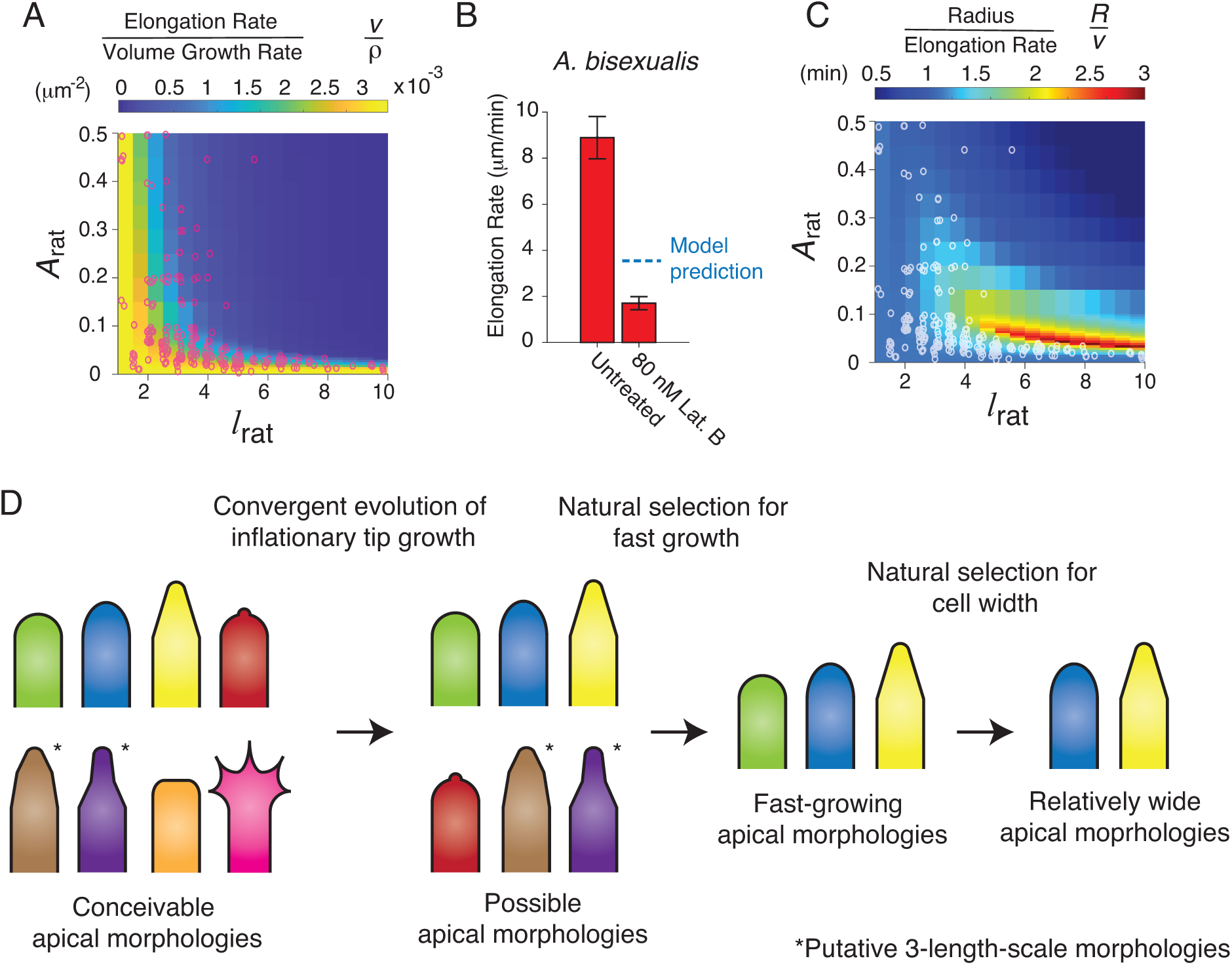
The morphospace of tip-growing cells corresponds to a fitness landscape. (A) The ratio of elongation rate, *v*, and total surface area expansion rate, *α*, versus *l*_rat_ and *A*_rat_ for simulated cells. The red circles indicate the same co-ordinates that yielded the best fit of the experimental morphologies shown in Fig. 5E,F. (B) The population-averaged experimental elongation rate of untreated *A. bisexualis* hyphae and those treated with 80 nM latrunculin B. Error bars indicate *±*1 s.d. *n* = 11 and 9 hyphae for untreated and treated cells, respectively. (C) The ratio of cell radius and elongation rate versus *l*_rat_ and *A*_rat_ for simulated cells. (D) Illustration of the evolutionary and developmental constraints on the apical morphologies of tip growing cells. (*) Denotes apical morphologies that are presumed to be possible given a 3-length-scale surface extensibility profile.

### Tapered morphologies balance the benefits of elongation rate and cell radius

Within the limited region of the morphospace that yields fast tip growth, it is still possible that the variation in apical morphology reflects selection for specific functions unrelated to growth rate. For example, it has been proposed that tapered morphologies promote invasive growth during pathogenesis [45], but this intuition has not been tested. We explored an alternate possibility: that tapered morphologies optimize a developmental variable, rather than an ecological one. In this light, one simple benefit of being tapered may be that it allows hyphae to increase their width beyond that prescribed by the primary length scale of cell-wall expansion. Although we discovered that cell widening leads intrinsically to reduced elongation rate (Fig. 6A,7A), we hypothesized that gradual sub-polar widening could be worth this reduction since fungi and oomycetes transport cytoplasm over long distances via dissipative flow [46], the energetic cost of which will be inversely related to cell radius. To explore this hypothesis, we calculated the cell radius per unit elongation rate, *R/v*, across our theoretical morphospace; this quantity describes how wide a cell is for a given elongation rate. Remarkably, *R/v* was maximal near the bistable region of the morphospace (Fig. 7C), meaning that cells with tapered morphologies not only grew relatively fast, but were also relatively wide for a rapidly elongating cell. That is, growing as a tapered cell is a way of simultaneously balancing *v/ρ* and *R/v*, which are explicit examples of complex traits.

## Discussion

We demonstrated that the convergent mechanical mechanism of inflationary tip growth intrinsically possesses an instability that imposes a strong empirical constraint on the natural apical morphologies of diverse tip-growing cells. Because cellular elongation rate is directly dependent on the same parameters as morphology, and because natural morphologies elongate faster than hypothetical morphologies, we conclude that the morphospace corresponds to a fitness landscape, rationalizing the constraint. Finally, competition between elongation rate and radius explains morphological variation within the natural fast-growing range of morphologies.

Thus, our analysis revealed three nested determinants of tip-growing morphology (Fig. 7D). First, inflationary growth imposes a mechanistic “developmental constraint” [47] on the apical morphology of tip-growing cells in the sense that morphologies not represented by our theoretical morphospace (Fig. 6B) could only be generated by adopting a completely different mechanism of cellular morphogenesis, for example, one where the cytoskeleton generates force. However, the effective pressure that the cytoskeleton can exert is approximately one hundred times smaller than than the typical turgor pressure of walled cells [48]. As a result, inflationary growth naturally results from the presence of turgor pressure and the cell wall, which may be viewed as the primary developmental constraints.

Due to the directionless nature of turgor pressure, any surface extensibility profile that decays over a single length scale will generate a rounded cell apex (Fig. S5C). The discovery that natural tip-growing cells employ surface extensibility profiles that decay over one or two length scales was empirical and inherently constrains apical morphologies. How would adding a third length scale affect morphology? Given our analysis, we propose that this would create the possibility of cell apices with “graduated” taper (*i.e.* where there are two sub-polar conical sections with different slopes; Fig. 7D). Although these morphologies are possible, we propose that they are still essentially just tapered cells, and therefore offer no advantage over those generated with a two-length-scale surface extensibility profile. If so, one- and two-length scale surface extensibility profiles generate the entire space of “useful” morphological variation.

Our most sweeping discovery was that an instability in inflationary tip growth imposes a second, strong constraint on morphology. Although the cusp bifurcation arises mechanistically from inflationary growth, the fact that it is a constraint was, *prima facie*, empirical. Tip-growing cells are ubiquitous across nature, serve essential functions in agriculture, are the source of many antibiotics, and are important pathogens of humans, crops, and fish. The finding of a widespread constraint on their morphology may have implications in any of these domains.

Given the constraint imposed by the cusp bifurcation, highly tapered apical morphologies are nearly unstable and therefore are vulnerable to perturbations (Fig. 6). This may explain why relatively few tip-growing species display tapered morphology and implicitly suggests that taper may serve an important function. One proposed function in oomycetes is that the tapered hypha acts as a “knife” during invasive pathogenesis [49]. This analogy breaks down when comparing the mechanics of a knife and a cell: while mechanical stress applied by a knife increases with increasing sharpness (i.e. taper), the mechanical stress applied by the apex of a tip-growing cell is simply equal to the turgor pressure, regardless of the taper.

We discovered an alternative function of taper, which imposed a third constraint: that it yields the widest possible rapidly elongating cells (Fig. 7C). The mycelia of many fungi and oomycetes grow as connected networks of hyphae in which cytoplasm moves via gross mass transport over macroscopic distances, a process that is energetically expensive for thin hyphae [46]. Interestingly, oomycetes exhibit a large variation in hyphal radii, and therefore taper [32]. Based on our analysis, we predict that highly tapered cells act as arterial hyphae that supply less tapered hyphal branches with cytoplasm. This will be an interesting subject for future research. The global constraint on apical cell morphology imposed by the cusp bifurcation only makes sense if we interpret the dependence of elongation rate on the morphospace as a fitness landscape (Fig. 7A). In other words, the assertion that fast tip growth is selected for by nature is a major inferential result of our analysis. On one hand, this is a simple conclusion that makes sense in terms of the ecology of these cells. For example, fungal, oomycete, and bacterial hyphae are the vegetative cells that compete within and between species to locate and consume nutrients. On the other hand, this a profound result that provides a window into the ecology - and rationalizes the morphology - of an enormous diversity of interesting and important species. It is well understood that instabilities can dramatically affect the dynamics of ecological systems [50, 51]. Similar dynamics in evolutionary systems have been predicted to exist theoretically [52]. Here, we explicitly demonstrate that an intrinsic instability in a fitness landscape can dictate evolutionary outcomes, thereby experimentally demonstrating a new conceptual paradigm for evolutionary-developmental biology.

In the language of life-history theory, selection for fast growth at the expense of morphological variation implies that tip-growing cells have undergone “*r*-selection,” meaning that in order to capitalize on resource abundance morphology is primarily selected insofar as it is correlated with rapid growth [53]. Therefore, just as the “function” of the shape of a dolphin is largely to minimize hydrodynamic drag (an external force), the main function of the morphology of tip-growing cells is to maximize the effect of turgor pressure (an internal force). More broadly, the real question is not whether cell morphology or growth rate are selected for - these are both crucial, complex phenotypes that will be pulled by natural selection in various ways for various reasons. Rather, we set out to explicitly quantify how such complex phenotypes interact in the context of evolution. In the case of tip-growing cells, growth rate won the evolutionary tug-of-war, which still however left room for functional morphological variation, particularly with respect to selection for cell width.

Our analysis therefore provides a paradigmatic case study in how the interaction of physical principles with natural selection can sculpt biological form. Since turgor pressure is directionless, inflationary growth tends to generate rounded apexes for the same reason that balloons are roughly round. While establishing a cell-wall surface extensibility profile with two length scales allows for tapered apical morphologies, these morphologies are nearly unstable since it is difficult to create the straight meridians on the conical region of these cells in the presence of pressure. Natural selection acts on this widespread developmental constraint. Therefore, tip growth morphogenesis specifically depends on the direct interplay between a fundamental physical principle (minimization of surface energy) and a fundamental evolutionary principle (fast reproduction outcompetes slow reproduction). Morphology itself may be useful only insofar as it obeys these nested constraints. As D’Arcy Thompson predicted, “the manifestations of adaptation become part of a mechanical philosophy” in which “*la Nature agit toujours par les moyens les plus simples*,” but also “*chaque chose finit toujours par s’accommoder a son milieu*” [54].

## Acknowledgments

We thank Jacques Dumais for helpful discussions. This work was performed in part at the Aspen Center for Physics. ERR and MOE were supported by NSF-CAREER Grant 2047404. BDZ was supported by the Ministry of Education, Science and Technological Development of the Republic of Serbia (451?03-68/2022?14/200053) and the Deutsche Forschungsgemeinschaft (GA 173/13-1). We thank Alice F.-X. Liang, Joseph Sall, Chris Petzold and Jason Liang at NYU Microscopy Laboratory for assistance with electron microscopy. The Microscopy Lab is partially supported by NYU Cancer Center Support Grant NIH/NCI P30CA016087, and NIH S10 OD019974.

## Materials and Methods

### Strains and Media

The strains used in this study are listed in Table S1. All fungal and oomycete cultures were cultured on yeast-malt growth medium (0.3% yeast extract (m/v), 0.3% malt extract, 0.5% peptone, 1% dextrose, and 1% agarose). Phycomyces sporangiophores were grown on potato dextrose agar. Pollen tubes were grown in 15 mM MES, 1.6 mM H_3_BO_3_, 0.1 mM KCl, 7.5% sucrose (m/v), adjusted to pH 5.3 with 0.1 mM KOH [55]. *S. venezuelae* was cultured in ISP media. *C. glutamicum* was cultured in BHI media.

### Imaging and image analysis

For imaging of *L. longiflorum* and *C. japonica* pollen tubes, 1 hour prior to imaging freezedried pollen grains grains were germinated in a growth medium. A thin layer (*<*1 mm) of solid growth medium (1% low melting point agarose) was deposited onto the bottom of a custom-made chamber, consisting of a dental polymer gasket adhered to a cover glass. While the agarose was still in a molten state, pollen grains were partially embedded in the solid medium by rinsing it with the liquid suspension of grains. The slide chambers were then filled with liquid medium. Cells that emerged from grains affixed to the agarose in this way were likely to grow along the gel-liquid interface.

For imaging fungal and oomycete hyphae (except for *A. arbuscula*) dental-polymer slide chambers were prepared, as above, with 1% low melting point agarose yeast-malt growth medium. After the agarose solidified, the chamber was inoculated with a small block of agarose from the growing front of the mycelium in the propagation culture. The top of the slide chamber was then sealed with a cover slip. After incubation for 48-72 hours at room temperature, the top of the chamber was opened, and filed with liquid medium for imaging.

For imaging *A. arbuscula*, cultures were propagated in cover-glass bottomed Petri dishes (MakTek). The bottom of the dishes were covered in a layer of *≈*2 mm of solid growth medium (1% agarose) and one edge of this substrate was inoculated. The hyphae grew in the thin layer of liquid between the cover glass and the agarose at the bottom of the dish. Cells were imaged 5-6 days after inoculation.

For imaging of *S. venezuelae*, 5 *µ*L of exponential phase culture was spotted onto a 1% agarose pad that was molded in a FastWell reagent barrier (Grace Biolabs).

For imaging of *C. glutamicum*, exponential phase cells were loaded into a bacterial microfluidic plate (CellASIC) and perfused with BHI media during imaging using the ONIX microfluidic system.

Time lapse images of pollen tubes, fungal and protistan hyphae, and bacterial cells were taken using an Eclipse TI2 inverted microscope (Nikon) and a Prime BSI sCMOS digital camera (Photometrics).

For imaging of *P. blakesleeanus*, a glass vial filled with solid media was inoculated with a heat-shocked spore suspension, and placed on custom stage equipped with 3-D mechanical adjustment and illuminated with an LED light (Tungsram) from above in a temperature-controlled room (22 *^◦^*C). Time-lapse images were taken using a digital CMOS camera (MicroQ) attached to the ocular tube of a horizontal stereo microscope (Leitz) [56].

All cell tracking was performed using custom MATLAB scripts. All software used in this study is available upon request.

Cell wall geometry for *S. pombe* and *M. truncatula* root hairs were obtained from previously published data sets [26, 57].

### Single-cell geometry and principal components analysis

To extract single-cell apical cell-wall geometry, we calculated the meridional curvature profile *κ*_s_ = *dφ*(*s*)*/ds* of the cell outline (i.e., cross-sectional meridian) extracted from each time point of a time-lapse image sequence (Fig. 2A,C-D). We then reconstructed the time-averaged apical geometry for each cell by integrating the time-averaged meridional curvature profile:

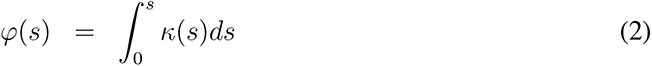

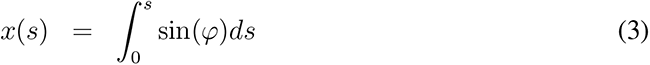

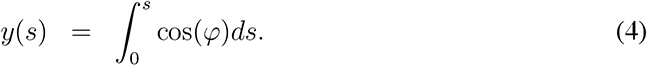

So as to only perform PCA on the apical region and not the distal region, we truncated the outlines using *φ* = [*−*0.95*π/*2, 0.95*π/*2] as bounds, and then interpolated the outlines with 75 equally spaced fiducial points. We found that *φ* = 0.95*π/*2 was the largest cutoff that did not include large portions of the cylindrical region of the cell. The axis of the apex was found by averaging opposing fiducial points on opposite sides of the cell meridian, and the outlines were rotated and translated such that their axes were aligned and that the cell pole was located at the origin. Principal components analysis was then applied to the interpolated (*x, y*) outlines at these points using the MATLAB command *pca*.

### Microsphere imaging and measurement of meridional speed

The meridional speed and expansion rate measurements for *Lilium longiflorum* pollen tubes (Fig. 2E,3G,J, S2E) was published previously [28]. For microsphere experiments related to *A. bisexualis* and *A. arbuscula*, 0.1 *µ*m aminated fluorescent microspheres (Sigma-Aldrich) were included in all media (10*^−^*^4^ dilution from stock), including solid media. For *A. bisexualis*, just after the slide chamber was filled with microsphere-containing media prior to imaging, the microspheres had a high affinity for the cell walls and bound to them. As the charges on the microspheres were neutralized, they lost their affinity for the cell wall, making it necessary to re-immerse them in freshly mixed microsphere growth medium every 10-15 minutes during imaging.

For *A. arbuscula*, a section of the solid media several millimeters in front of the leading edge of the mycelium was excised with a scalpel, forming a well to which microsphere-containing liquid media was added. Over the course of minutes, the microspheres diffused through the liquid layer between the cover glass as the solid media and reached the mycelium, whereupon they adhered to the hyphal cell walls.

For *L. longiflorum* pollen tubes, to adhere the microspheres to cells, a syringe pump was used to load a glass microcapillary (inner diameter *≈* 50 *µ*m) with microsphere-containing media. The microcapillary was mounted on a micromanipulator and was used to address elongating pollen tubes. Pressure was applied manually to the syringe pump to stream microspheres past the growing pollen tubes during growth. After 5-10 seconds of streaming the suspension past the tip, the exterior of the cell was effectively coated in microspheres. The microsphere medium was continually streamed past the cells during imaging such that their cell walls were constantly covered as they expanded.

Fluorescent microsphere tracking was performed using custom MATLAB scripts [28]. From the tracks of the microspheres (Fig. S2A) and the cell surface geometry, we calculated the meridional speed of the microspheres (Fig. S2B-D), which is the speed at which the microspheres move along the cell meridian in the frame of reference of the cell pole. To calculate the principal expansion-rate profiles from the experimental measurements of meridional speed, the spatial dependence of the meridional speed was fit with the empirical function *v*(*φ*) = *v*_0_ sin *φ*(1 + *c*_1_*φ* + *c*_2_*φ*^2^ + *· · ·*) [35]. F-tests were used to determine the maximum number of free parameters, *c_i_*, that could be retained in this function without fitting noise in the data. The principal expansion rates were calculated from the time-averaged cell geometry profile and this fit of the meridional speed profile using the equations:

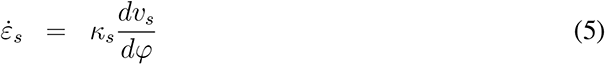

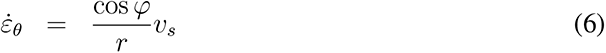

where *r*(*s*) is the local radius, or the radial distance of a point on the cell surface from the cell axis (Fig. 2A).

Because the meridional speed and the cell wall geometry specifically determine the principal expansion rates, to fit these spatially dependent variables together is to also fit the principal expansion rates. The free fit of the meridional speed corresponds to the fit of the fully anisotropic mechanical model (Eq. 16, below) shown in Fig. S2C-E. To fit the experimental meridional speed and cell-wall geometry given isotropical mechanical properties (Fig. 2C- E), we followed a method that we previously developed [58]. Note that in previous analyses, isotropical mechanical properties have been distinguished from transverse (in-plane) mechanical properties where the cell walls is stratified in the normal dimension. However, since we used a two-dimensional formulation of cell surface mechanics, there is no difference between these cases.

The principal tensions were calculated from the experimental profiles of the meridional and azimuthal curvatures, using Eqs. 7-10 below. An arbitrary pressure of 1 atm was assumed. This choice does not affect the spatial distributions of the tensions.

### Simulations of cell growth and parameter space search

Simulations of tip growth with arbitrary expansion rates or extensibility profile were performed with a custom MATLAB routine. An initial hemispherical geometry was defined with *N* (an odd number typically between 101 and 301) discrete (*x, y*) co-ordinates. The arclength, *s*, was calculated on the geometry, with the polar point defined as *s* = 0 (Fig. 2A). The surface extensibility (or expansion-rate) profile(s) to be simulated was then prescribed on these points. The radius of the hemisphere was arbitrary but was generally chosen to be somewhat larger than the largest length scale associated with the extensibility profile.

When the surface extensibility profile was prescribed, at each step of the simulation, first the principal curvatures in the cell wall were calculated [30]:

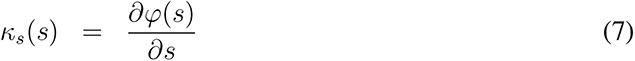

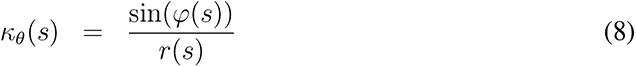

where *r*(*s*) is the radial distance from the cell axis to the fiducial point. From the principal curvatures, the principal tensionss were calculated using using force-balance equations [35]:

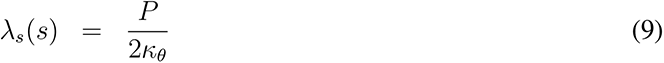

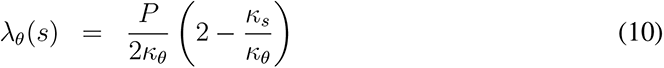

In general, the turgor pressure was arbitrarily defined since it only affects the global magnitude of the expansion rates, and therefore the rate of tip growth, rather than the cellular morphology. Unless principal expansion rate profiles were prescribed, they were calculated rom the principal tensions using Eq. 1 of the main text:

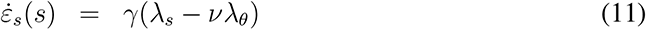

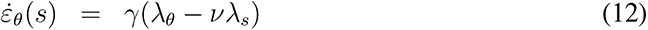

To then calculate the velocity of each fiducial point we inverted the kinematic equations [30]:

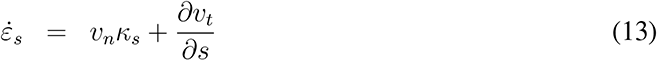

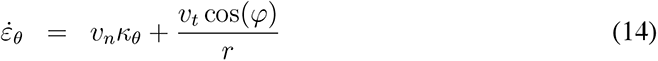

where *v_n_* and *v_t_* are the velocity components of the discrete points in the directions normal and tangential to the cell outline. Finally, the displacement of each point *v_n_dt*, *v_t_dt* was calculated (where *dt* was the time between steps), converted to cartesian co-ordinates, and added to the cell geometry to find the updated cell geometry. This updated geometry was smoothed with a smoothing spline (the MATLAB *csaps* command, with smoothing factor 0.9999), which was critical to prevent small errors from causing runaway instabilities. This routine was iterated until cell geometry converged. At each step, the time step was updated to ensure that the displacement at the pole converged to a fixed, small value.

To validate the simulation routine, we input the experimental surface extensibility profiles, which were calculated to perfectly fit the experimental apical morphology [58]. In our simulations, this surface extensibility profile yielded a meridional curvature profile that was indistinguishable from the experimental profile (Fig. S5).

For the parameter space searches described in Fig. 5 and Fig. S7, *l*_rat_ was varied on the domain (1, 10) at 0.25 increments and *A*_rat_ was varied on the domain (0, 0.1) at 0.005 increments and the domain (0.1, 0.5) at 0.05 increments. For each value of *A*_rat_, the search was initiated at *l*_rat_ = 1 and *A*_rat_ was increased incrementally to 0.5, using the solution of cell shape from the previous parameter co-ordinate as the initial condition for each simulation.

To resolve the fold bifurcation shown in Fig. 6B, we focused on the domain *l*_rat_ *∈* (1, 10) (increments of 0.25) and *A*_rat_ *∈* (0, 0.1) (increments of 0.0025). We searched the space as before, but both increased and decreased *A*_rat_ for a given *l*_rat_. To resolve the fold bifurcation we found (*l*_rat_*, A*_rat_) co-ordinates where there were two stable solutions. For each one of these co-ordinates, from the two stable apical geometries, 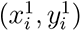 and 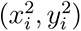, we found ten intermediate geometries 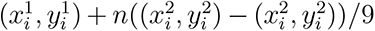, where *n* = 0, 1*, …,* 9. We then initiated a simulation with each of these geometries and determined to which stable solution the simulation converged (Fig. S7F). An unstable solution was computed to be the mean of neighboring geometries that converged to different stable fixed geometries (dotted line, Fig. S7F). From these unstable geometries, the radius could be calculated to calculate the whole manifold of solutions shown in Fig. 6B.

To calculate the best fit of the model for each experimental cell geometry, we first found the time-averaged cell outline, which was found by integrating the time-averaged meridional curvature profile using Equations 2-4. We discretized the time-averaged experimental cell outline into the same number of points that was used to simulate tip growth in the computational parameter space search. We then calculated the normalized average error,

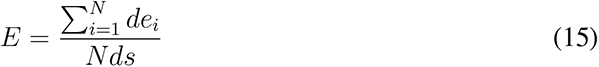

which is the average error of each discretized point relative to the interpoint arclength. We found values of *l*_rat_ and *A*_rat_ that minimized this error.

The normalized elongation rate (Fig. 7A) was calculated as follows. At steady-state, the total rate at which the cell increases in volume is *ρ* = *πR*^2^*v*, where *R* is the cell radius and *v* is the elongation rate of the cell. Therefore, a normalized form of the elongation rate is 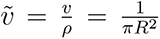. This quantity has dimensions of inverse area: it is the rate at which the cell elongates per unit volume added to the cell. We arbitrarily dimensionalized the morphospace in Fig. 7A using the radius of a pollen tube. This specific dimensionalization does not affect the landscape of *v*.

The dependence of the ratio of cell radius and elongation rate was calculated by first calculating the absolute (un-normalized) radius and elongation rate of the simulations. We then dimensionalized these variables by arbitrarily assuming that cells that are generated by a single-length-scale surface extensibility profile grow with the radius and elongation rate of a pollen tube. This specific dimensionalization does not affect the landscape of *R/v*.

### Theoretical analysis of anisotropical version of the linear extensibility model

Equation 1 of the main text is a generic linear formulation of inflationary tip growth if the wall is isotropic within the plane of the cell wall, meaning that cell wall polymers are not synthesized in an oriented manner. In previous formulations of tip growth (e.g., [35]), “transverse” or “plane” isotropy was distinguished from isotropy when considering stratification of cell wall material in the normal direction. However, in our two-dimensional formulation isotropy and transverse isotropy are the same thing, and we therefore use the word isotropy to mean isotropic in the two-dimensional sense.

Although it is unlikely that a large degree of structural anisotropy (resulting form oriented synthesis) can develop across the apical cell wall during tip growth since the polar cell wall must be isotropic, it was important to demonstrate this or demonstrate that the possibility of structural anisotropy in the cell wall does not alter our results. Here, we perform two analyses that demonstrate both of these results.

i. First, the fully anisotropic version of Eq. 1 of the main text is the most general version of the linear extensibility implementation of the inflationary mechanism of cell growth:

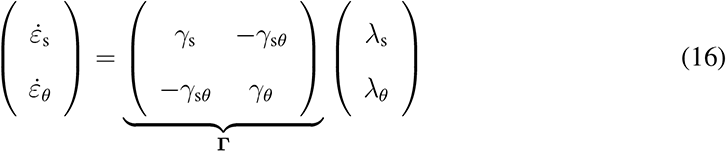

The diagonal elements of **Γ** (*γ*_s_ and *γ_θ_*) are the anisotropic surface extensibilities and determine how fast the wall expands in a given direction if uniaxial tension is applied in that direction. These quantities are equal at the cell pole due to rotational symmetry but may be unequal in the sub-polar cell wall if oriented cell wall polysaccharides were synthesized there. The offdiagonal element *γ*_s_*_θ_* mechanically couples the principal directions by determining how fast the wall contracts in one direction due to uniaxial tension in the orthogonal direction. This matrix equation can be re-written:

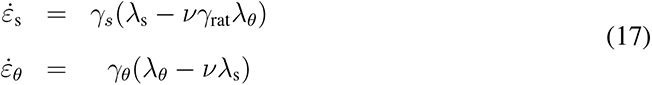

where here 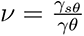 and 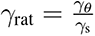 is the ratio between the principal surface extensibility profiles.

Solving for the surface extensibility profiles, we find:

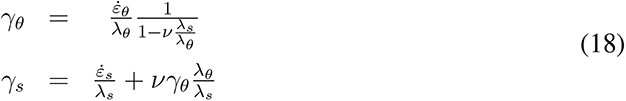

where 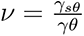 is the only quantity that we can not experimentally measure. Substituting experimental values for the principal expansion rate and tension profiles into Eq. 18 reveals that for all reasonable values of 0 *< ν* ¡ 0.75, the principal surface extensibility profiles are approximately equal across the apical cell wall (Fig. S3), indicating that the degree of anisotropy in the cell wall is negligible. Further allowing for *ν* to be spatially inhomogeneous allows the profiles to be precisely equal (theory not shown).

ii. Second, note that it is clear that the azimuthal surface extensibility profiles for *A. bisexualis* and *A. arbuscula* (and probably their meridional profiles) decay over two length scales, regardless of anisotropy or the flow coupling, *ν* (Fig. S3A,B). That is, two length scales in the azimuthal extensibility profile is directly implied by the kinematics of tip growth of these species, independent of whether any anisotropy is present.

This result allowed us to perform a parameter space search similar to the one we performed for the isotropic case, but for the case where anisotropy develops due to the meridional surface extensibility profile decaying with one length scale. The key morphological parameters in this case are the ratio of the two length scales associated with the azimuthal surface extensibility profile and the ratio of the amplitudes of the two component profiles. We explore the case where the meridional surface extensibility profile decayed with a single length scale equal to the smaller length scale of the azimuthal profile (Fig. S4A), generating anisotropy. This parameter space search gave rise to a morphospace that was very similar to the morphospace generated by the isotropic version of the model (Fig. S4B). Furthermore, this morphospace possessed a cusp bifurcation that constrained natural apical morphologies in the exact same manner as that from the isotropical model (Fig. S4C,D). In other words, the cusp catastrophe is a generic feature of inflationary tip growth that constrains the apical morphologies of tip growing cells and that does not depend on the specific molecular architecture of the cell wall. Therefore, although our data strongly suggest that the cell wall is not structurally anisotropic (Fig. S3), they also implies that even if anisotropy were to develop, it does not alter any of our results.

Next, we explicitly demonstrated that structural anisotropy alone cannot explain non-rounded apical morphologies if the azimuthal surface extensibility does not decay with two distinct length scales. To do this, we simulated tip growth where the cell wall developed structural anisotropy sub-polarly by imposing that the two principal surface extensibility profiles decayed with distinct length scales, but where each one decayed over a single length scale. To simulate azimuthal structural anisotropy (*γ_θ_ > γ*_s_), we imposed that the azimuthal surface extensibility profile was a Gaussian that decayed over a single length scale, *l* = *A e*^(*−s/l_θ_*)2^ (Fig. S4E). To obtain the meridional surface extensibility profile, we multiplied this first profile by a spatially dependent extensibility ratio, 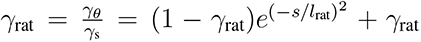 that also decayed with a single length scale, *l*_rat_ and where *γ*_rat_ is the maximum degree of anisotropy (Fig. S4E). In this case, the key morphological parameters are the ratio *l*_rat_*/l_θ_* and *γ*_rat_. A parameter space search across all possible morphological parameters revealed that no combination of parameters could produce highly tapered apical morphologies characteristic of *A. bisexualis* or accurately explain the curvature profile of *A. arbuscula* (Fig. S4F). Indeed, nearly the entire parameter space yielded rounded shapes with values of P1 ≲ 3, whereas the majority of natural species have higher values of P1 (Fig. 2F). A parallel analysis in which meridional structural anisotropy (*γ*_s_ *< γ_θ_*) developed sub-polarly revealed similar results (Fig. S4G,H).

Finally, the anisotropical version of inflationary growth (Eq. 16) provides negligible additional fitting power over the isotropical version (Eq. 1; Fig. 2C-E, see above for fitting strategy).

To summarize our analysis of tip growth with structural anisotropy, analysis of the expansion rate profiles strongly suggest that the cell wall of *A. bisexualis* and *A. arbuscula* are not anisotropic (Fig. S3). However, even if weak anisotropy were to develop sub-polarly, the azimuthal surface extensibility would have to decay with two length scales, which would lead to a cusp bifurcation in the morphospace that constrains natural morphologies, quantitatively similar to that observed for the isotropic case (Fig. S4).

### Fluorescence cell wall labeling of *A. bisexualis*

To fluorescently label the cell walls of *A. bisexualis* hyphae, we cultured them on cover-glass-bottomed slides, as above. Calcofluor white (10 *µ*M; Sigma-Aldrich) or Direct Red23 (0.1 mg/mL, Sigma-Aldrich) were dissolved in YMM media. To control for binding kinetics of the dye, we also included 1 M sorbitol in the media, which plasmolyzes the hyphae, leaving just the cell wall. We added the dye/sorbitol media to growing hyphae and incubated fro 30 min, and then imaged. The fluorescence intensity along the cell wall was analyzed with custom MATLAB algorithms.

### Transmission electron microscopy of *A. bisexualis*

*A. bisexualis* hyphae were fixed with 2.5% glutaraldehyde in 0.075M phosphate buffer (pH 7.2) for 30 mins at room temperature, and then at overnight at 4 *^◦^*C. The samples were washed 3 times with 0.075M phosphate buffer (10 minutes each time), post-fixed with 1% osmium tetroxide for 1.5 hour at room temperature, and dehydrated in the serial of ethanol solutions (30, 50, 70, 85, 95, 100%; 10 min each). The hyphae were then washed in acetone two times for 10 min each, infiltrated with acetone and embedded in Spurr (Electron Microscopy Sciences). Semi-thin sections (500 nm) were stained with 1% toluidine blue. Ultrathin sections (75 nm) were cut and mount on a formvar coated slot grid and stained with uranyl acetate and lead citrate by standard methods. Stained grids were imaged with Gemini300 scanning electron microscope (Zeiss) using the STEM detector at 20 kV with working distance of 3.3mm and scan speed of 8, the image pixel size is 5.6 nm.

The cell wall geometry and exocytic vesicles from TEM micrographs were extracted manually. Vesicles were binned by distance from the cell pole along the cell axis, and then mapped to the point on the cell outline at the same axial distance. The vesicles density at each point along the cell outline was then calculated as the total number of vesicles at that point divided by 2*πrdl*, where *dl* is the length of a bin and *r* is the radius at that point.

### Latrunculin B treatment of cells

Latrunculin B (abcam) was dissolved in DMSO to make a 2.5 mM master stock solution, which was further diluted 1000 fold in water to make 2.5 *µ*M Latrunculin B stock solution. This was used to prepare yeast malt media containing 1% agarose and 80 nM latrunculin B. Molten media was poured as a thin (*≈* 3*mm*) layer into petri dishes and allowed to solidify. A circular punch was used to cut out sections of this agarose. These sections were then placed on top of *A. bisexualis* hyphae that were growing in cover glass imaging chambers described above. An additional 100 *µ*L of 80nM Latrunculin B solution was added to the top of the agar pad . The mycelia were left to grow overnight at room temperature before imaging.

To calculate the theoretical prediction of the effect of latrunculin B on cell elongation rate shown in Fig. 7B, we used the expression *ρ* = *πR*^2^*v*, where *ρ* is the rate of volume increase. For the untreated case we calculated *ρ* by substituting experimental values of *R* and *v*, and found the predicted value of *v* for the latrunculin-treated cells by substituting in their mean radius, *R*, and the calculated value of *ρ*.

