## Supplemental Figures for "A fitness landscape instability governs the morphological diversity of tip-growing cells"

### Supplementary Figure Legends

**Fig. S1. Apical geometry of tip-growing cells varies across the tree of life.** (A) Phase micrographs of nine tip-growing organisms. (B) Representative time-averaged meridional curvature (normalized by cell radius) versus arclength (normalized by radius) for cells from the species shown in (A). (C) Percent of morphological variation across the entire multi-species data set explained by the first five principal components. (D) Length of the cell apex versus the first principal variable. (E) Taper of the cell apex versus the first principal variable. (F) Meridional curvature (normalized by cell radius) versus arclength (normalized by radius) for four values of P1 corresponding to the population-averaged values of four tip-growing species. For this calculation, the remaining principal variables were fixed to be the average experimental value across all cells from all species.

**Fig. S2. Inflationary tip growth explains the mechanics of cell-wall expansion.** (A) The tracks of fluorescent microspheres mapped onto the location of the cell wall during *A. bisexualis* tip growth. (B) The kinematic variables of tip growth. (C-E) Meridional speed versus arclength for *A. bisexualis*, *A. arbuscula*, and *L. longiflorum*. The profiles are representative of $n = 2, 4, 4$ cells, respectively. The orange lines are the best fit with the linear model of tip growth given by Eq. 1 of the main text. Fitting the meridional speed is equivalent to fitting the principal strain rates since the latter are calculated from the former (see (B), Eq. 5,6). Next to each profile is the sum of squared residuals (SSR) for the best fits from the isotropic (Eq. 1) and anisotropic (Eq. 16) implementations of inflationary growth, demonstrating that the anisotropic version of the model does not increase fitting power over the isotropic version. (F) The experimental principal expansion rate profiles for an *A. bisexualis* hypha, exhibiting azimuthal anisotropy. (G) A simulation of tip growth inputting the principal expansion rates shown in (F). (H) A pair of principal expansion rate profiles exhibiting isotropy found by enforcing that the two profiles both equal the mean of the experimental ones shown in (F). (I) A simulation of tip growth inputting the principal expansion rates shown in (H). (J) A pair of principal expansion rate profiles exhibiting meridional isotropy found by enforcing that the meridional expansion-rate profile equal the experimental azimuthal profile, and vice versa (i.e., by "switching" the profiles). (K) A simulation of tip growth inputting the principal expansion rates shown in (J). (L) Taper from simulations of tip growth for azimuthal anisotropy, isotropy, and meridional anisotropy, derived from experimental profiles of *A. bisexualis*, *A. arbuscula*, and *L. longiflorum* cells. (M) Schematic of structural anisotropy.

**Fig. S3. The cell wall is structurally isotropic**. This grid shows the principal surface extensibility profiles that are predicted from the experimental principal surface expansion rates and principal surface tensions, calculated using Eq. 18. Profiles from one cell from each of our model systems are shown. Because if we allow for anisotropy, we cannot constrain the flow coupling, we calculate what the principal surface extensibility profiles would be, given a range of values of this parameter. This analysis demonstrates that regardless of the flow coupling, the principal surface extensibilities must be approximately isotropic to generate the experimental surface extensibility profiles. Not shown are cases where the predicted principal surface extensibilities have regions of negative values, which is unphysical (dotted boxes).

**Fig. S4. Structural anisotropy within the cell wall does not explain morphological diversity** (A) Empirical anisotropic principal surface extensibility profiles. The azimuthal surface extensibility profile is a Gaussian mixture and therefore decays with two length scales, whereas the meridional profile decays with a single length scale equal to the shorter of the two length scales associated with the azimuthal profile. This creates structural anisotropy. (B) The theoretical morphospace generated by linear inflationary tip growth combined with the empirical surface extensibility profile shown in (A). P1 is the first principal component from our multispecies PCA analysis (Fig. 2F). A flow coupling of 0.5 was used. The morphospace qualitatively and quantitatively resembles the morphospace for the isotropic case (Fig. 5B). (C) Normalized radius as a function of $A_{\text{rat}}$ for $l_{\text{rat}} = 9$. This slice through the solution manifold is indicated by the green line in Fig. (B). $A_{\text{rat}}$ was cycled to demonstrate hysteresis. (D) Coordinates of $(l_{\text{rat}}, A_{\text{rat}})$ that, when used to simulate cell growth, yielded the best fit of the geometry for each cell of each species (white circles) overlaid onto the morphospace shown in (B). (E) Illustration of the strategy to explore the effect of azimuthal structural anisotropy on apical morphology without imposing two length scales on either of the principal surface expansion rate profiles. (left) A Gaussian azimuthal surface extensibility profile, $\gamma_\theta(s)$, that decays with a single length scale, $l_\theta$ was defined. (center) This profile was multiplied by a spatially dependent Gaussian anisotropy ratio, $\gamma_{\text{rat}}(s) = \gamma_\theta/\gamma_{\text{s}}$, that also decayed with a single length scale that was less than $l_\theta$. (right) This produced principal surface extensibility profiles that possessed azimuthal anisotropy and where each decayed with a single length scale. (F) The morphospace generated by a parameter space search across the two morphogenetic parameters associated with the principal expansion rates shown in (E) can only produce round cells (with low values of P1). (G) Illustration of the strategy to explore the effect of meridional structural anisotropy on apical morphology. (left) A Gaussian meridional surface extensibility profile, $\gamma_{\text{s}}(s)$, that decays with a single length scale, $l_{\text{s}}$ was defined. (center) This profile was multiplied by a spatially dependent Gaussian anisotropy ratio, $\gamma_{\text{rat}}^{-1}(s) = \gamma_{\text{s}}/\gamma_\theta$, that also decayed with a single length scale that was less than $l_{\text{s}}$. (right) This produced principal surface extensibility profiles that possessed meridional anisotropy where each decayed with a single length scale. (H) The morphospace generated by a parameter space search across the two morphogenetic parameters associated with the principal expansion rates shown in (G) can only produce round cells (with low values of P1).

**Fig. S5. The Gaussian mixture surface extensibility profile predicts apical geometry**. (A-C) i) Experimental time-averaged apical geometry and meridional curvature versus arclength. ii) The apical geometry and meridional curvature generated simulations of tip growth inputting the experimentally derived surface extensibility profiles (Fig. 4A-C). iii) The apical geometry and meridional curvature generated by simulations of tip growth inputting the experimental Gaussian mixture function (Fig. 4F) that provided the best fit of the experimentally derived surface extensibility profiles. iv) The apical geometry and meridional curvature obtained from simulations of tip growth inputting a function of the form $\gamma(s) = A_1 \cos^2(s/l_1) + A_2 \cos^2(s/l_2)$ that provided the best fit of the experimentally derived surface extensibility profiles. For (C) iv), the apical geometry and meridional curvature were obtained from simulations of tip growth inputting a single Gaussian that provided the best fit of the experimentally derived surface extensibility profile of *L. longiflorum.*

**Fig. S6. The biochemical composition of the cell wall does not change across the cell apex**

(A) The spatial probability distribution of vesicle density versus arclength extracted from TEM images. Confidence intervals indicate $\pm$ 1 s.d. $n = 3$ cells. (B) Representative micrograph of a cell apex stained with Direct Red 23. (C) Population-averaged fluorescence intensity versus normalized arclength of Direct Red 23-labeled cells. Confidence intervals indicate $\pm$ 1 s.d. $n = 17$ cells. (D) Representative micrograph of a cell apex stained with Calcofluor White. (E) Population-averaged fluorescence intensity versus normalized arclength of calcofluor white-labeled cells. Confidence intervals indicate $\pm$ 1 s.d. $n = 15$ cells. (F) Population-averaged normalized meridional curvature versus normalized arclength. The pink shaded area indicates the primary length scale associated with the decay of the curvature profile, and therefore also the smaller length scale associated the surface-extensibility profile.

**Fig. S7. The flow coupling does not influence global morphological variation.** (A) Time-averaged apical geometry for an *A. bisexualis* cell (blue line) and the apical geometry of a cell simulated using Eq. 1 of main text combined with a Gaussian mixture surface extensibility profile with $l_{\text{rat}}$ and $A_{\text{rat}}$ that yielded the best fit of the experimental morphology (red line). (right) The error associated with the best fit is calculated for each point used to discretize the outlines. (B) The distribution of mean error (averaged across points for a given cell) between the experimental outlines and the best fits provided the theoretical morphospaces generated by inflationary growth with a two-length-scale surface extensibility profile. The four distributions correspond to the best fits provided by morphospaces generated using 4 values of flow coupling, demonstrating that the flow coupling does not influence the fitting power of the model. (C) P1 of simulated cells using Eq. 1 of the main text and a Gaussian mixture surface extensibility profile, as a function of $l_{\text{rat}}$ and $A_{\text{rat}}$, for four different values of $\nu$. (D) Apical geometries of simulated cells for four different values of $\nu$ at a two points in the morphospace shown in (C). (E) Normalized radius as a function of $A_{\text{rat}}$ for $l_{\text{rat}} = 9.5$. This slice through the solution manifold is indicated by the red line in Fig. 6B of the main text. The dotted section of the curve represents unstable solutions. (F) Values of $(l_{\text{rat}}, A_{\text{rat}})$ that, when used to simulate cell growth, yielded the best fit of the geometry for each cell of each species overlaid onto contours of normalized radius, and showing the empirical constraint by the bistable region of the morphospace. (G) The population-averaged radius of untreated *A. bisexualis* hyphae and those treated with 80 nM latrunculin B. Error bars indicate $\pm 1$ s.d. $n = 11$ and 9 hyphae for untreated and treated cells, respectively.

### Supplementary Figures

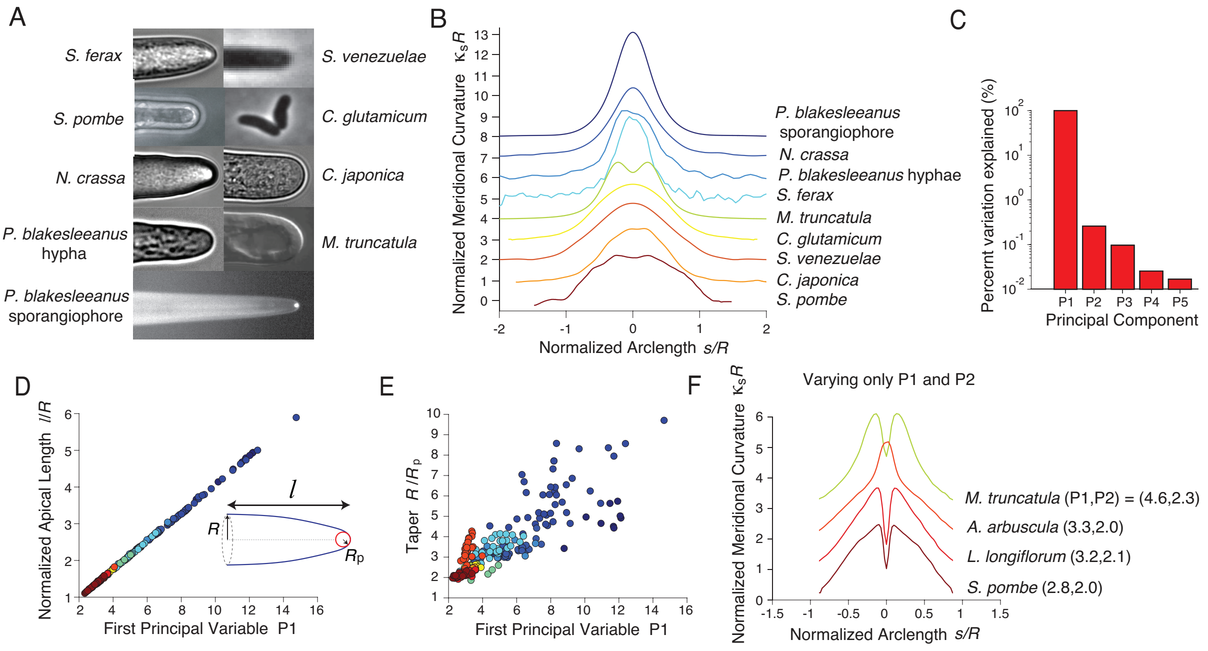


**Figure S1**

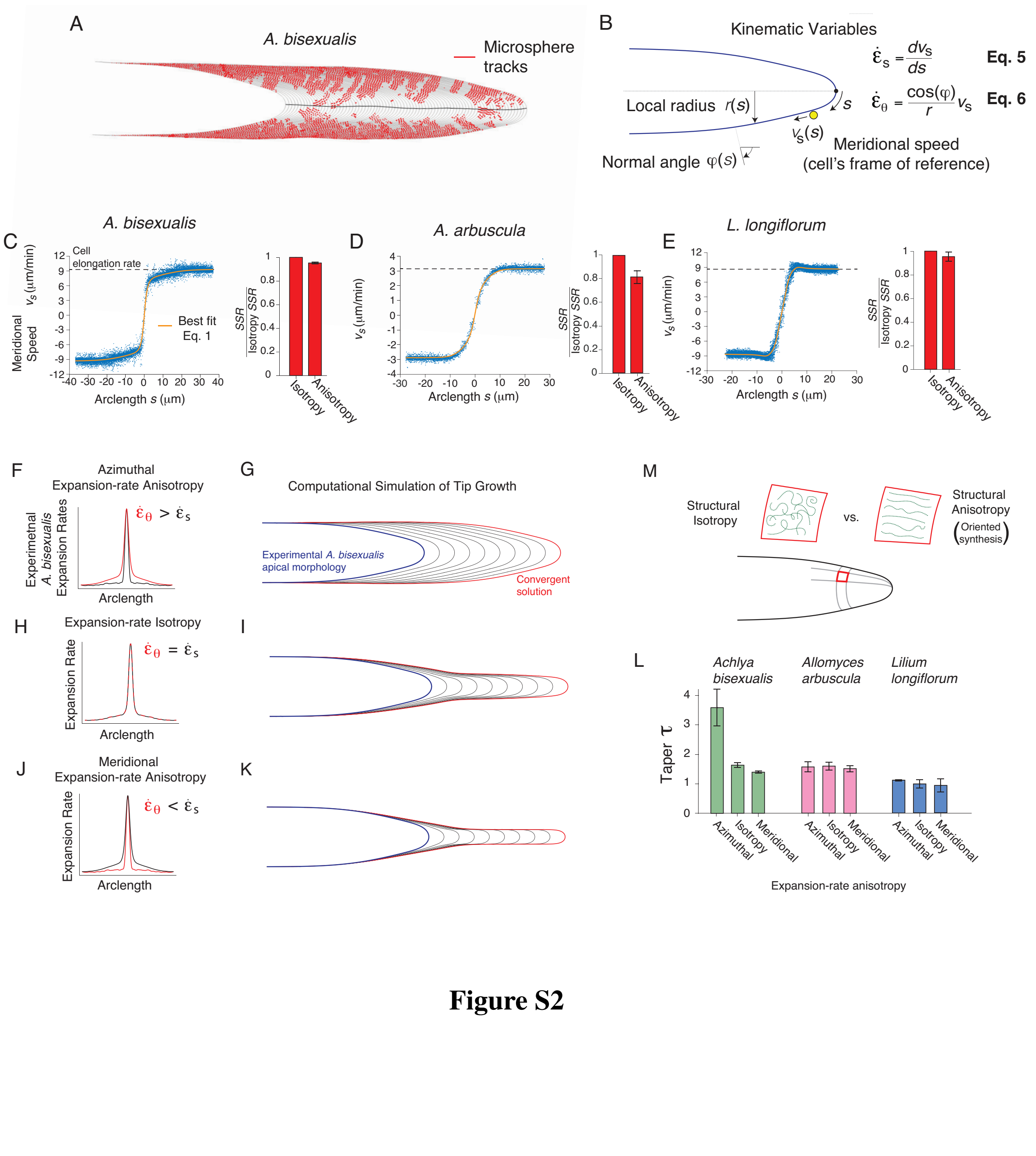

A
A. bisexualis
Microsphere tracks
B
Kinematic Variables
Local radius r(s)
Normal angle φ(s)
$v_s(s)$
Meridional speed (cell's frame of reference)
$\dot{\varepsilon}_s = \frac{dv_s}{ds}$ Eq. 5
$\dot{\varepsilon}_\theta = \frac{\cos(\varphi)}{r} v_s$ Eq. 6
C
A. bisexualis
Cell elongation rate
Best fit Eq. 1
Meridional Speed $v_s$ (μm/min)
Arclength s (μm)
SSR / Isotropy SSR
Isotropy
Anisotropy
D
A. arbuscula
E
L. longiflorum
F
Azimuthal Expansion-rate Anisotropy
Experimental A. bisexualis Expansion Rates
$\dot{\varepsilon}_\theta > \dot{\varepsilon}_s$
Arclength
G
Computational Simulation of Tip Growth
Experimental A. bisexualis apical morphology
Convergent solution
H
Expansion-rate Isotropy
Expansion Rate
$\dot{\varepsilon}_\theta = \dot{\varepsilon}_s$
I
J
Meridional Expansion-rate Anisotropy
$\dot{\varepsilon}_\theta < \dot{\varepsilon}_s$
K
M
Structural Isotropy
vs.
Structural Anisotropy
(Oriented synthesis)
L
Achlya bisexualis
Allomyces arbuscula
Lilium longiflorum
Taper τ
Azimuthal
Isotropy
Meridional
Expansion-rate anisotropy


**Figure S2**

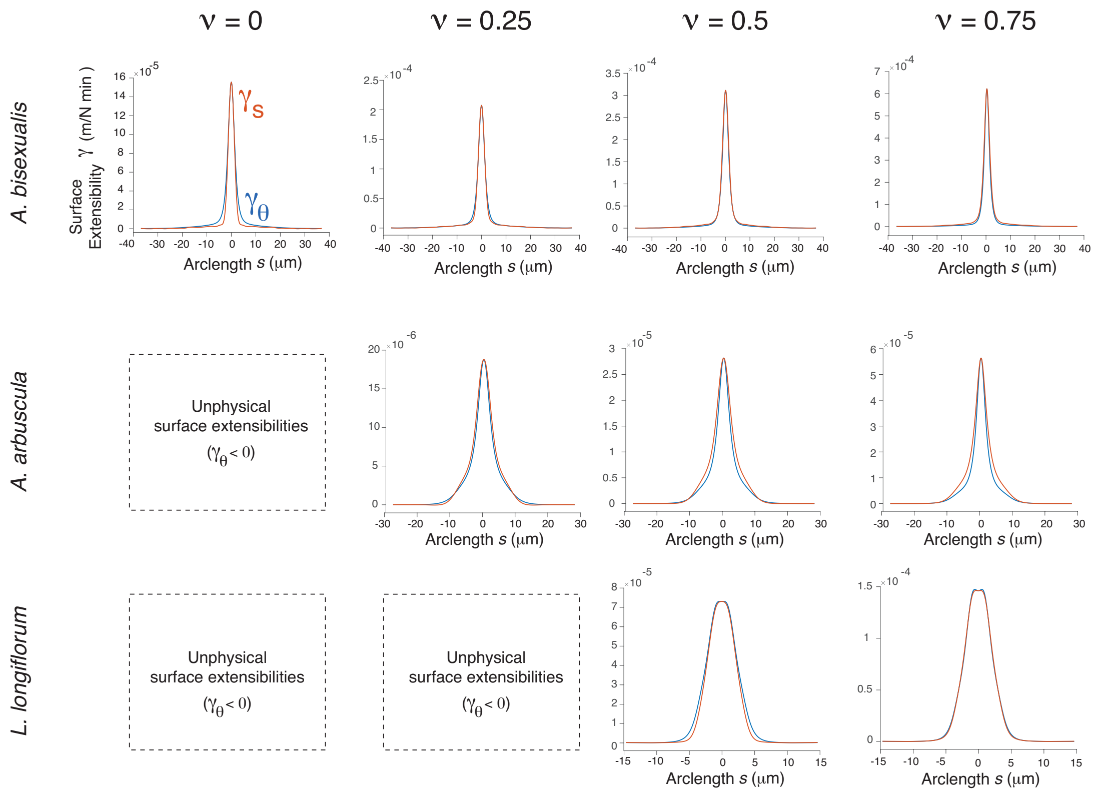
ν = 0
ν = 0.25
ν = 0.5
ν = 0.75
A. bisexualis
A. arbuscula
L. longiflorum
Surface Extensibility γ (m/N min)
γs
γθ
Arclength s (μm)
Unphysical surface extensibilities (γθ < 0)

**Figure S3**

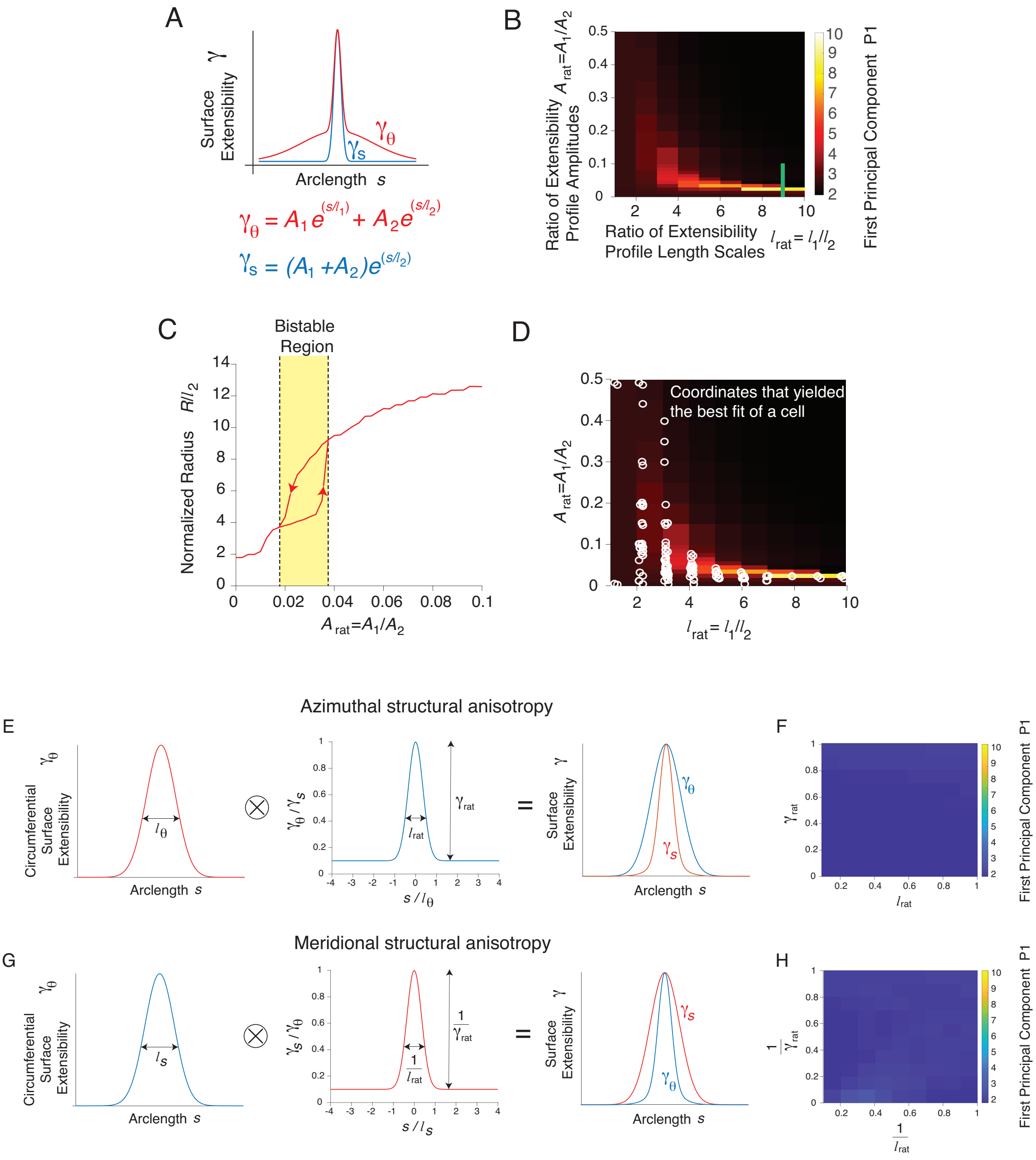
A
Surface Extensibility γ
Arclength s
$\gamma_\theta = A_1 e^{(s/l_1)} + A_2 e^{(s/l_2)}$
$\gamma_s = (A_1 + A_2) e^{(s/l_2)}$
B
Ratio of Extensibility Profile Amplitudes $A_{rat}=A_1/A_2$
Ratio of Extensibility Profile Length Scales $l_{rat}=l_1/l_2$
First Principal Component P1
C
Bistable Region
Normalized Radius $R/l_2$
$A_{rat}=A_1/A_2$
D
Coordinates that yielded the best fit of a cell
$A_{rat}=A_1/A_2$
$l_{rat}=l_1/l_2$
Azimuthal structural anisotropy
E
Circumferential Surface Extensibility $\gamma_\theta$
Arclength s
$\gamma_\theta / \gamma_s$
$s / l_\theta$
Surface Extensibility γ
F
$\gamma_{rat}$
$l_{rat}$
First Principal Component P1
Meridional structural anisotropy
G
Circumferential Surface Extensibility $\gamma_\theta$
Arclength s
$\gamma_s / \gamma_\theta$
$s / l_s$
Surface Extensibility γ
H
$\frac{1}{\gamma_{rat}}$
$\frac{1}{l_{rat}}$
First Principal Component P1

**Figure S4**

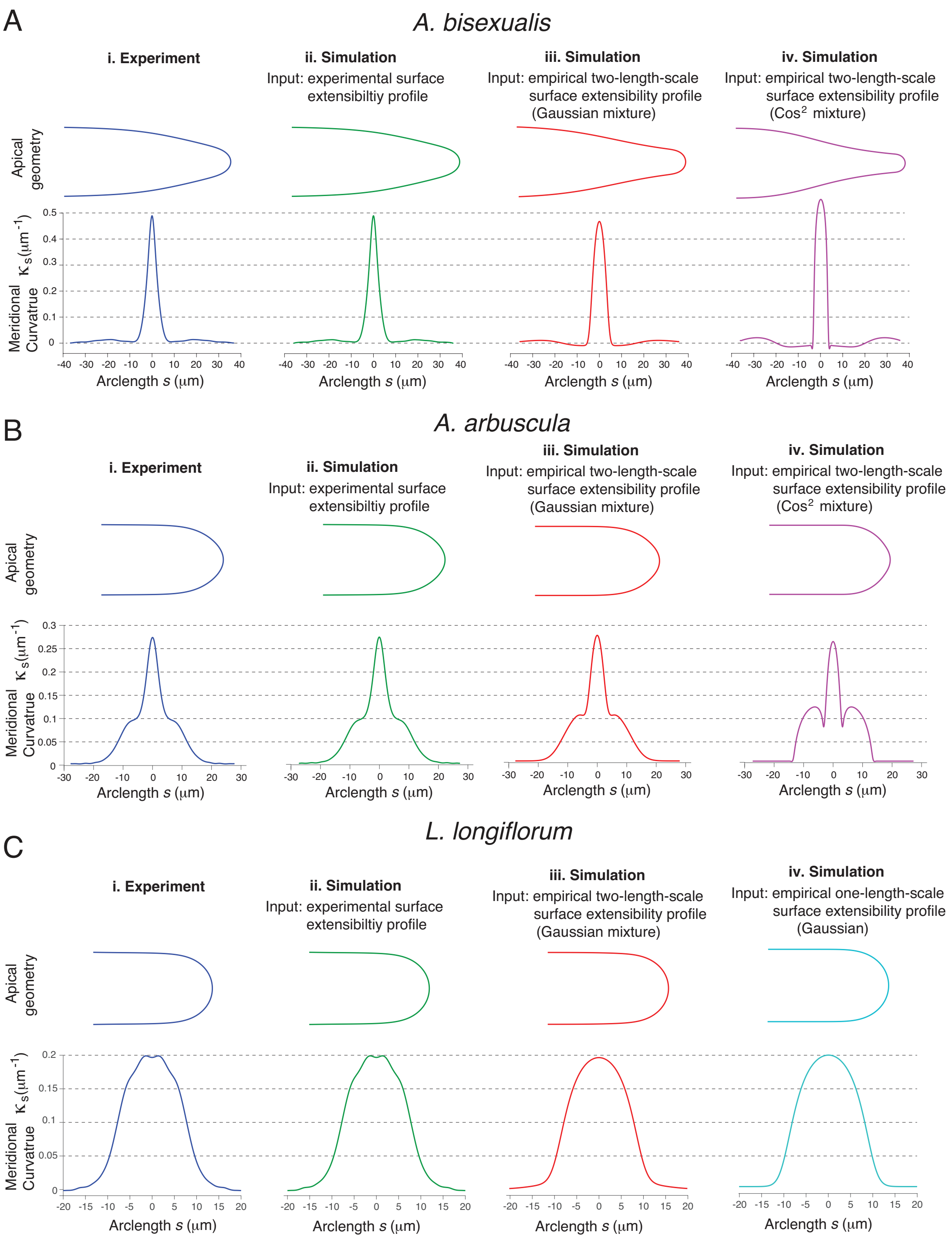
A
A. bisexualis
i. Experiment
ii. Simulation
Input: experimental surface extensibiltiy profile
iii. Simulation
Input: empirical two-length-scale surface extensibility profile (Gaussian mixture)
iv. Simulation
Input: empirical two-length-scale surface extensibility profile (Cos² mixture)
Apical geometry
Meridional Curvatrue κs(μm⁻¹)
Arclength s (μm)
B
A. arbuscula
i. Experiment
ii. Simulation
Input: experimental surface extensibiltiy profile
iii. Simulation
Input: empirical two-length-scale surface extensibility profile (Gaussian mixture)
iv. Simulation
Input: empirical two-length-scale surface extensibility profile (Cos² mixture)
Apical geometry
Meridional Curvatrue κs(μm⁻¹)
Arclength s (μm)
C
L. longiflorum
i. Experiment
ii. Simulation
Input: experimental surface extensibiltiy profile
iii. Simulation
Input: empirical two-length-scale surface extensibility profile (Gaussian mixture)
iv. Simulation
Input: empirical one-length-scale surface extensibility profile (Gaussian)
Apical geometry
Meridional Curvatrue κs(μm⁻¹)
Arclength s (μm)

**Figure S5**

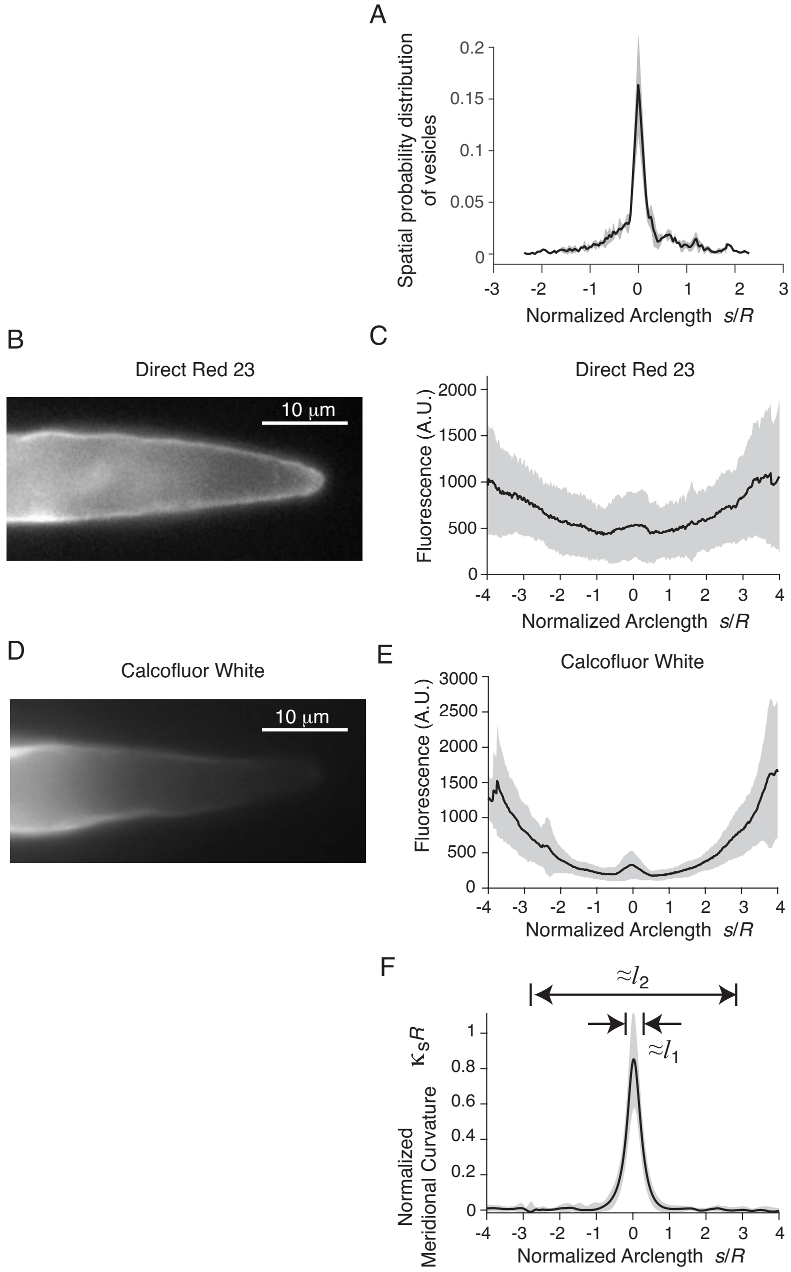
A
Spatial probability distribution of vesicles
Normalized Arclength s/R
B
Direct Red 23
10 μm
C
Direct Red 23
Fluorescence (A.U.)
Normalized Arclength s/R
D
Calcofluor White
10 μm
E
Calcofluor White
Fluorescence (A.U.)
Normalized Arclength s/R
F
≈l2
≈l1
κsR
Normalized Meridional Curvature
Normalized Arclength s/R

**Figure S6**

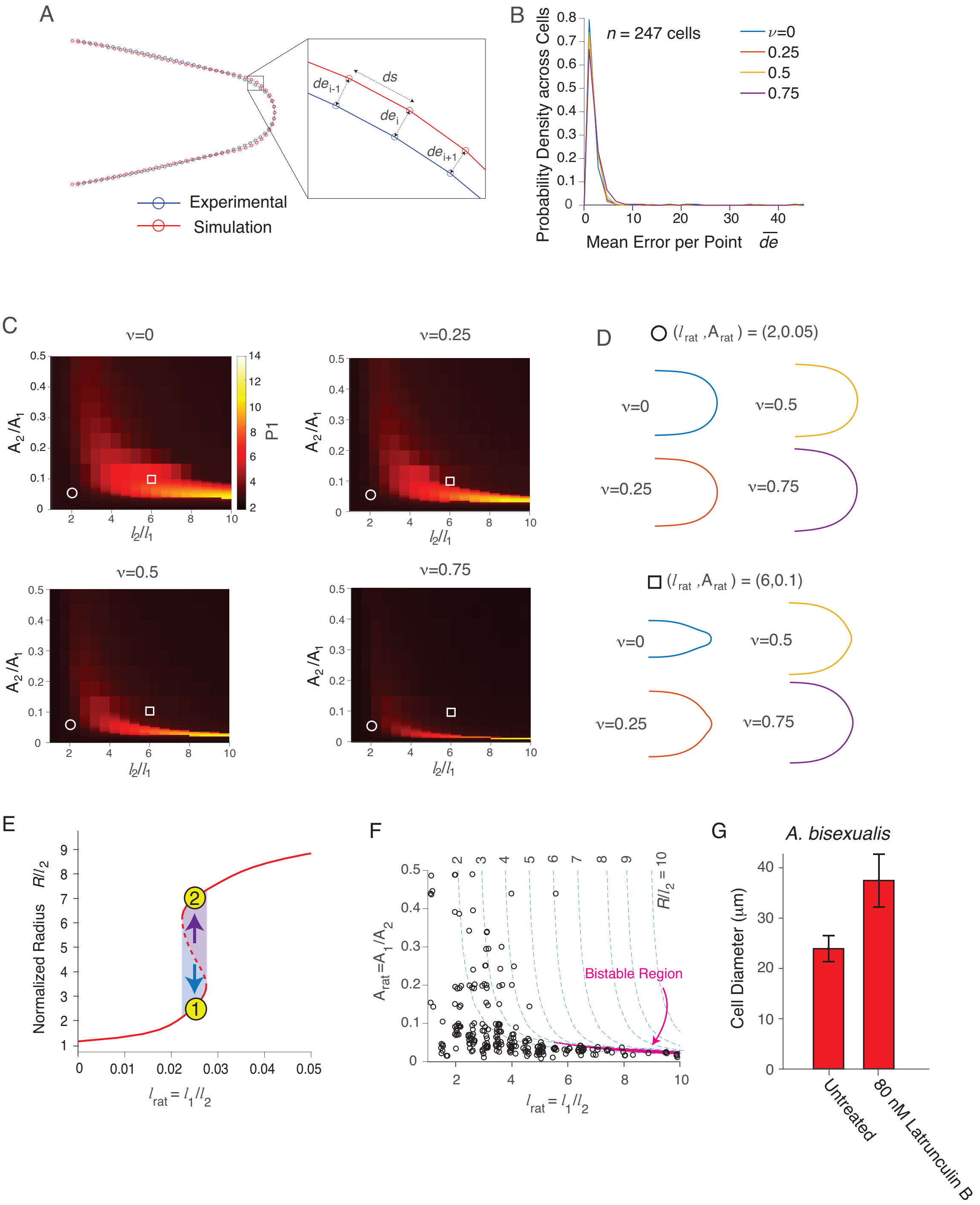
A
Experimental
Simulation
ds
$de_{i-1}$
$de_i$
$de_{i+1}$
B
n = 247 cells
ν=0
0.25
0.5
0.75
Probability Density across Cells
Mean Error per Point $\overline{de}$
C
ν=0
ν=0.25
ν=0.5
ν=0.75
$A_2/A_1$
$l_2/l_1$
P1
D
$(l_{rat}, A_{rat})$ = (2,0.05)
$(l_{rat}, A_{rat})$ = (6,0.1)
E
Normalized Radius $R/l_2$
$l_{rat} = l_1/l_2$
F
$A_{rat}=A_1/A_2$
$R/l_2$ = 10
Bistable Region
$l_{rat} = l_1/l_2$
G
A. bisexualis
Cell Diameter (μm)
Untreated
80 nM Latrunculin B

**Figure S7**
